# A Rapid *in vivo* Pipeline to Identify Small Molecule Inhibitors of Amyloid Aggregation

**DOI:** 10.1101/2022.04.11.487937

**Authors:** Muntasir Kamal, Jessica Knox, Robert I. Horne, Om Shanker Tiwari, Andrew R. Burns, Duhyun Han, Dana Laor Bar-Yosef, Ehud Gazit, Michele Vendruscolo, Peter J. Roy

## Abstract

Amyloids are associated with over 50 human diseases and have inspired significant effort to identify small molecule remedies. Here, we present a novel *in vivo* platform that efficiently yields small molecule disruptors of amyloid formation. We previously identified small molecules that kill the nematode *C. elegans* by forming membrane-piercing crystals in the pharynx cuticle, which is rich in amyloid-like material. We show here that many of these molecules are known amyloid-binders whose crystal-formation in the pharynx can be blocked by amyloid-binding dyes. Furthermore, we found that amyloid fibrils can seed small molecule crystal formation *in vitro*. These observations suggest that small molecule crystals are seeded by the cuticle’s amyloid-like material. We asked whether this phenomenon could be exploited to identify additional molecules that interfere with the ability of amyloids to seed higher-order structures. We screened 2560 compounds and identified 85 crystal suppressors, which we found to be 10-fold enriched in known amyloid disruptors relative to a random set. Of the uncharacterized suppressors, we found 25% to inhibit Ab42 fibril nucleation and/or extension *in vitro*, which is a hit rate that far exceeds other screening methodologies. Hence, screens for suppressors of crystal formation can efficiently reveal small molecules with amyloid-disrupting potential.

## Introduction

Previous small molecule screens with the free-living nematode *C. elegans* serendipitously revealed compounds that visibly accumulate within the heads of worms ^1–4^. This accumulation can be observed using a low magnification dissection light microscope ^5^. We demonstrated that the accumulated small molecules form either birefringent irregular objects, globular non-birefringent spheres, or a mixture of the two, depending on the structure of the compound ^5^. Birefringence is an optical property that reveals the subject to harbour repeating units that uniformly alter the angle of incidence of polarized transmitted light. We showed that these birefringent objects: i) are observable within minutes of exposure of the worms to the small molecule; ii) contain the exogenous small molecule; iii) grow over time; iv) are formed exclusively in association with the pharynx cuticle; and v) are not due to the consumption of precipitate in the media by the worm ^5,6^. Given their birefringent nature and growth over time, we refer to the birefringent objects as crystals for simplicity’s sake, although their exact biophysical nature remains to be determined (Figure 1A-C). The crystals kill younger animals likely because they perforate the plasma membrane of surrounding cells and rapidly grow to occlude the pharynx lumen ^5^.

**Figure 1.**
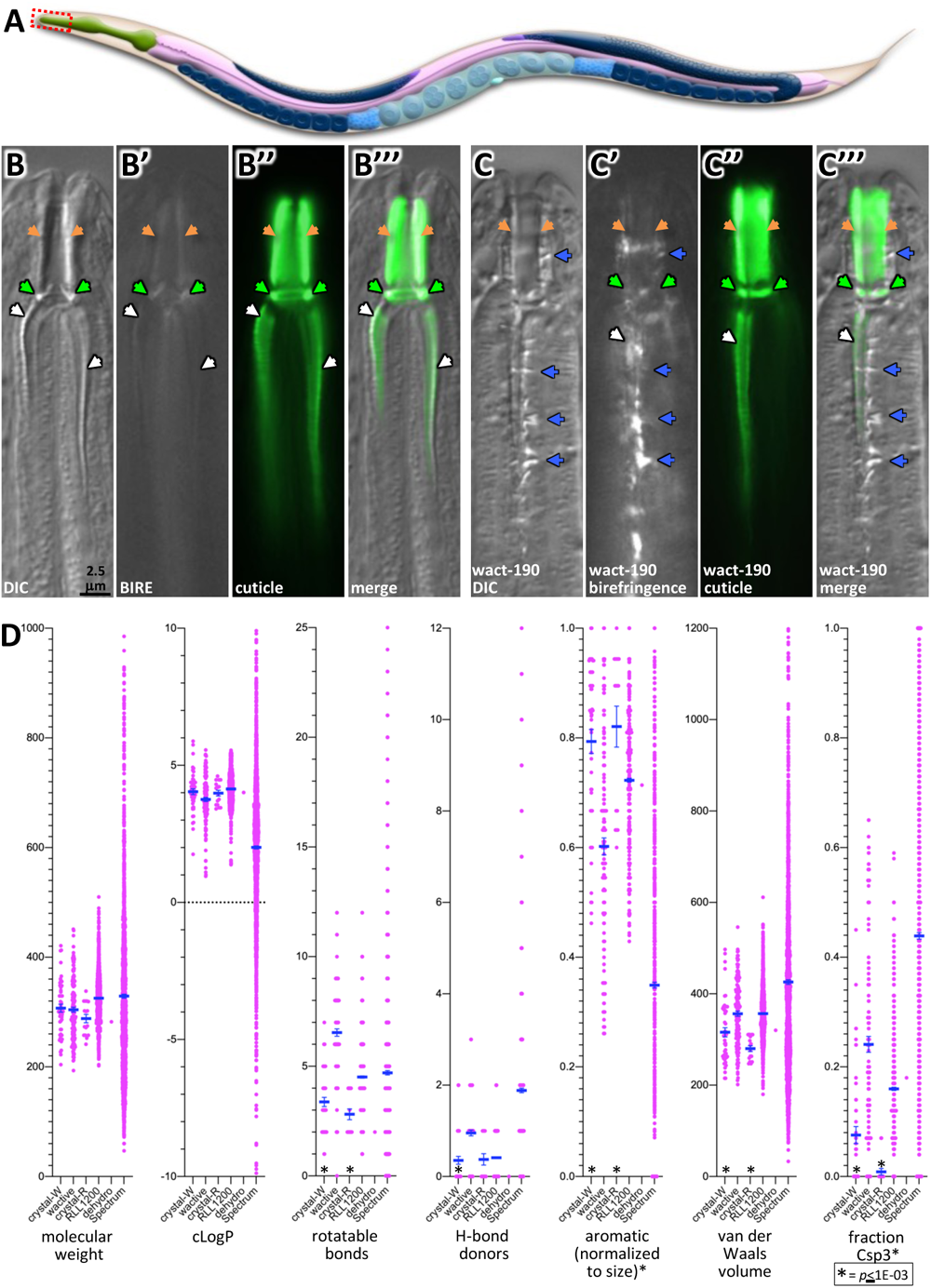
Crystal-Forming Small Molecules are Compact and Aromatic. **A.** A schematic of *C. elegans* anatomy (from WormAtlas; used with permission). The pharynx is green. **B & C.** Anterior-most region of a worm expressing the IDPC-1::GFP cuticle marker ^9^ treated with either 1% DMSO solvent control (B) or 30 μM wact-190 (C) for six hours. Orange arrows indicate the buccal cavity cuticle, the green arrows highlight the buccal collar, the white arrows indicate the cuticle that lines the channels, and the blue arrowheads highlight the birefringent crystals. Differential interference contrast (DIC), birefringent (BIRE), and the IDPC-1::GFP (cuticle) images are shown. The scale bar in B applies to all images in the upper right-hand side. **D.** Crystalizing molecules have significantly fewer rotatable bonds and hydrogen-bond donors, but greater aromaticity, and compactness (as measured by van der Waals volume and fraction Csp3) compared to non-crystalizing molecules from the same library. The mean and standard error is shown.

More recently, we described how the crystals form in tight association with the non-luminal face of the *C. elegans* pharyngeal cuticle ^6^. This cuticle is a flexible non-collagenous chitin-rich matrix that is repeatedly shed and rebuilt at each larval stage ^7,8^. We also constructed a spatiotemporal map for the development of the pharyngeal cuticle of *C. elegans* ^9^. That study revealed the pharynx cuticle to be highly enriched with intrinsically disordered proteins (IDPs) and proteins with intrinsically disordered regions (IDRs) ^9^. Many of these IDP and IDR-rich proteins have amyloidogenic features that include an enrichment in π-bonding and hydrogen-bonding capabilities, and have protofilament formation propensity ^9^. Furthermore, we and others showed that amyloid-binding dyes specifically stain the pharyngeal cuticle ^9,10^. Despite these findings, neither transmission electron microscopy (TEM) nor diagnostic tests provide evidence for extended amyloid fibrils within the pharyngeal cuticle ^9,11–13^, which is consistent with the flexible nature of the cuticle ^14,15^. Taken together, these observations indicated that the cuticle is rich in amyloid-like material that lacks the highly ordered nature of amyloid fibrils.

Here, we show that molecules that form crystals *in vivo* can indeed form a crystalline lattice *in vitro* through π-bond interactions. We go on to show that over 50% of the 29 distinct structural scaffolds that form crystals *in vivo* are reported amyloid binders. Consistent with these observations, we find that, i) nascent crystals distort the amyloid-like material within the cuticle without a concomitant effect on the chitin matrix; ii) Aβ42 fibrils can seed the formation of objects *in vitro* from a small molecule that forms crystals *in vivo*; and iii) worms incubated with the amyloid-binders Congo Red and Thioflavin S can block crystal formation. Collectively, these results suggest that the amyloid-like material in the pharynx cuticle can seed the formation of small molecule crystals.

Given that well-characterized amyloid binders can block the seeding of crystals in the pharynx, we wondered whether the crystallization phenomenon could be exploited to identify other small molecules that block the ability of amyloids to seed higher order structures such as amyloid fibrils. We therefore screened 2560 drugs and natural products for those that could suppress crystal formation in the *C. elegans* pharynx. The screen yielded 85 reproducible suppressors, several of which are fluorescent and specifically localize to the cuticle, which is consistent with their expected site of action.

The 85 crystal suppressors can be grouped into 45 distinct small molecule scaffolds. Of these 45, one third are previously characterized disruptors of amyloid fibril formation. We tested 44 of the suppressors that have not been previously implicated in amyloid disruption and found that 11 (25%) inhibit Aβ42 aggregation. In total, 40 of the 85 suppressors (47%) were shown to disrupt amyloid formation of one type or another, which is a hit rate higher than most previously reported screening methodologies ^16–18^. We conclude that screens for suppressors of crystal formation within the *C. elegans* pharynx cuticle can enrich for small molecules with amyloid-disrupting potential.

## Results

### Crystallizing Molecules are Aromatic, Compact, and Crystalize via π Bond Interactions

We previously identified 38 compounds that form crystals in association with the *C. elegans* pharynx cuticle by surveying 238 compounds from our custom library of <u>w</u>orm-<u>act</u>ive (wactive or wact) small molecules ^2,5^. We have since identified additional crystalizing molecules from a targeted survey of the wactive library molecules to yield a total of 48 crystallizing compounds (Supplemental Figure 1).

To better understand how these 48 crystallizing molecules differ from the 149 true negatives that fail to crystalize, we extended our previous physicochemical analysis to analyze a total of 72 features (Supplemental Table 1). We found no major differences in molecular weight (*p*=0.75) or hydrophobicity (i.e., cLogP) (*p*=0.05), and as previously reported ^5^, the crystallizing molecules have fewer rotatable bonds (*p*=4.3E-05) and fewer hydrogen-bond donors (*p*=2.4E-06) than non-crystalizing wactives (Figure 1D). Our more in-depth analysis shows that the crystallizing wactives are highly aromatic (*p*=2.6E-10) and more compact (i.e., smaller van der Waals volume, *p*=7.3E-04). A measure that combines these features, called fraction Csp3, reports the ratio of sp3 hybridized carbon atoms over the total carbon atom count of the small molecule ^19^ and reveals the compactness of the crystallizing molecules (*p*=5.5E-09).

To test whether these features are specific to crystallizing molecules from the wactive library, we screened two other libraries for crystallizing molecules. The first library was a custom collection of 1178 small molecules called RLL1200 (Supplementary Datafile 1). The second was the Spectrum library of 2560 drugs and natural products (Microsource Inc). To identify crystalizing molecules from these libraries, we first screened them for those that kill L1 larvae in our standard 6-day viability assay ^2^, reasoning that crystalizing molecules invariably kill larvae ^5^ and would be among the hits. We identified 70 lethal molecules from the RLL1200 library and 78 from the Spectrum library. To find crystallizing molecules among these hits, we exploited our previous finding that the *sms-5(0)* null mutant resists crystal-induced death by blocking crystal formation ^5^. Re-screening the lethal hits against the *sms-5(0)* mutant revealed 16 molecules from RLL1200 and 1 molecule from Spectrum that: i) kill wild type; ii) fail to kill *sms-5(0)* mutants; and iii) form crystals in association with the pharynx (Supplementary Datafile 1). The physicochemical features of the RLL1200 crystallizing molecules are not significantly different from those of the wactive crystallizing molecules (*p*>0.001) (Figure 1D; Supplemental Table 1; Supplementary Datafile 1). Statistical analysis was not conducted with the one Spectrum hit, but its properties are similar to the means of the other crystalizing molecules. Hence, compact aromatic properties are common to most crystallizing molecules.

Given that aromatic structures are rich in π bonds, we asked whether π bond interactions might play a role in the crystallization of the small molecules *in vitro*. To investigate, we used crystal powder-X-ray diffraction (PXRD) with two exemplar molecules wact-190 and wact-416. Wact-190 crystallized in the triclinic crystal system with the space group P-1, whereas wact-416 crystallized in the orthorhombic crystal system with the space group Pna2_1_ (Figure 2A-C, 2F-H; Supplemental Table 2; Supplemental Figure 2). The centroids of the aromatic rings for the respective crystalized molecules had distances of 5.0 Å or less, indicating that π-π interactions play a key role in their assembly into higher ordered structures ^20^ (Figure 2D and I). We compared the crystallization results with their simulated data to validate the bulk phase purity of the crystals. In comparison to the computer-simulated data, the small molecule crystals displayed a similar crystalline nature, peak-pattern matching and single-phase identification, confirming the individual structural purity of the samples (Figure 2E and J; Supplemental Figure 2).

**Figure 2.**
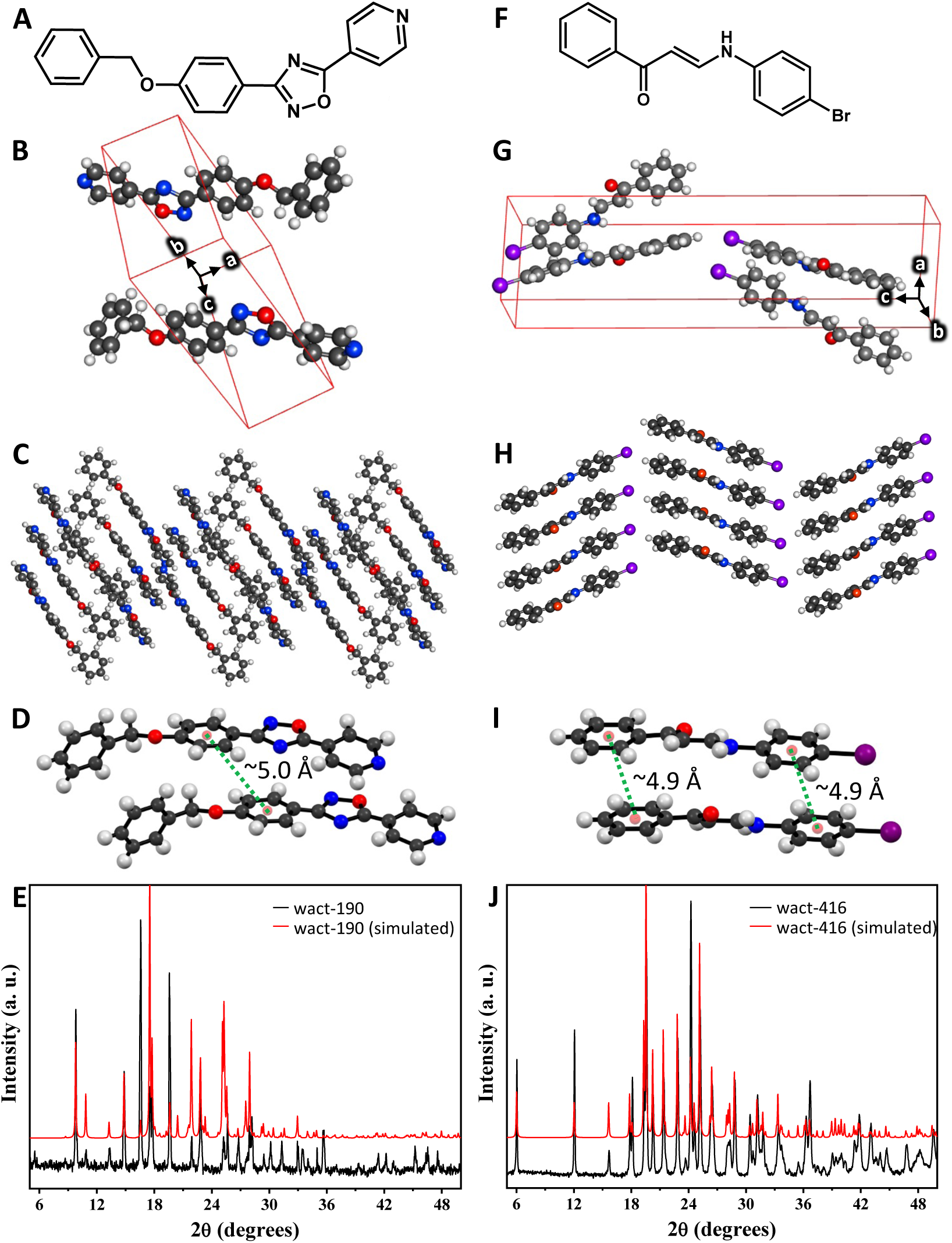
Biophysical Analyses of *in vitro* Small Molecule Crystals. **A.** The chemical structure of wact-190. **B.** The unit cell with wact-190 molecules whose centroid lies within the unit cell with the three coordinates shown with white letters and black corresponding arrows. The wact-190 crystal unit cell shows two molecules per asymmetric unit cell in a triclinic and anti-parallel packing arrangement. **C.** The 3D crystal packing arrangement along the b-axis (see Supplemental Figure 2a-2c for other aspects). **D.** The calculated distance between the indicated centroid (pink sphere) of two adjacent molecules stacked along crystallographic b-axis within the wact-190 crystal is approximately 5.0 Å. **E.** The experimental and simulated PXRD patterns of wact-190 molecule (see Supplemental Figure 2d for more details). **F.** The chemical structure of wact-416. **G.** The unit cell with wact-416 molecules whose centroid lies within the crystal unit cell. The wact-416 crystal unit cell shows four molecules per asymmetric unit cell in an orthorhombic and parallel packing arrangement. **H.** The 3D crystal packing (view along crystallographic b-axis) shows parallel packing for wac-416 crystals (see Supplemental Figure 2e-2g for other aspects). **I.** The calculated distance between the indicated centroid (pink sphere) of the 6-membered rings of two adjacent molecules stacked along crystallographic b-axis within the wact-416 crystal is approximately 4.9 Å. **J.** The experimental and simulated PXRD patterns of wact-416 molecule (see Supplemental Figure 2h for more details).

### Molecules that Induce Crystal Formation in *C. elegans* are Enriched with Amyloid-Binding Substructures

To investigate the structural diversity among crystal-forming compounds, we created a structural-similarity network by first reducing each molecule to its core scaffold. We then linked the scaffolds that had a Tanimoto similarity coefficient of 0.8 or more (see methods). This produced a network of 6 clusters that are unconnected with one another and 15 singletons (Figure 3A). Visual inspection of the molecules within the network reveals 30 structurally distinct families (Figure 3A; Supplemental Figure 1).

**Figure 3.**
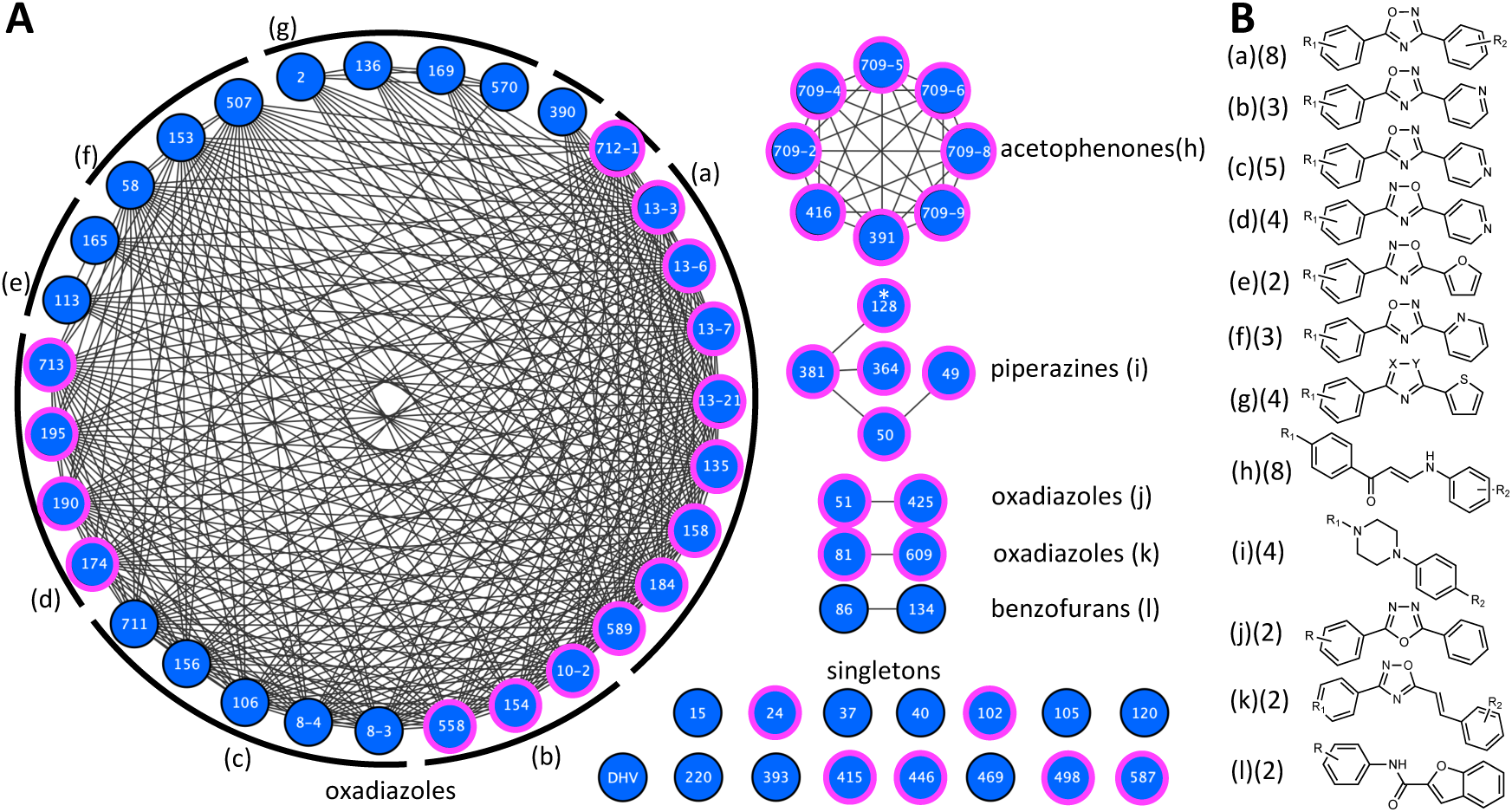
A Structural Similarity Network of 65 Crystal-Forming Compounds. **A.** An unsupervised structural similarity network of the 65 crystalizing molecules. Each node represents a small molecule, labelled with a wactive number or DHV (dehydrovariabilin). The link represents a Tanimoto score of structural similarity of the core scaffold of 0.8 or higher. The lower-case labels indicate groups of molecules with the same core scaffolds as indicated in B. Nodes highlighted in pink indicate a core scaffold known to interact with amyloids (see Supplemental Table 3 and Supplemental Data File 1 for details). The asterisks on the wact-128 node highlights that visual inspection reveals that it is a distinct scaffold despite its connection to wact-381 and represents the 30^th^ unique family. **B.** Markush representations of the core scaffolds of structural analogs able to crystalize in the worm. R, X, and Y groups are shown in Supplemental Figure 1.

A search of the patent and academic literature using the SciFinder search tool ^21^ revealed that 15 of the 30 scaffolds (50%) are found in molecules that bind amyloids (pink-outlined nodes in Figure 3A, Supplemental Figure 1, and Supplemental Table 3). For example, scaffolds (a), (k) and close analogs of wact-102 have been shown to bind Aβ42 amyloid aggregates with low nanomolar affinity ^22–24^. Scaffold (i) and close analogs of wact-128 and wact-415 were shown to bind prion amyloid aggregates with low nanomolar affinity ^25^. In another example, wact-498 is a flavonoid, which is an intensively studied class of molecules that have therapeutic potential for treating amyloid-related neurodegeneration (reviewed ^26^). In particular, isoflavone analogs of wact-498 can disrupt Aβ42 plaque formation in a *C. elegans* model of the human disease ^27^ and were shown to directly bind Aβ42 *in vitro* ^28^. The same analysis was performed on 62 randomly chosen scaffolds from molecules that do not form crystals ^5^. We found only 6 scaffolds (10%) that were previously described to bind amyloids (Supplemental Table 4). These data suggest that the collection of crystal-forming molecules is enriched with amyloid-associating substructures.

### Crystals Disrupt the Amyloid-Like Material of the Pharyngeal Cuticle

We sought to better understand how crystal growth perturbs the materials of the cuticle. To investigate, we used cuticle-binding dyes and fluorescent reporters. We previously found that the red-fluorescent Congo red (CR) and blue fluorescent Thioflavin S (ThS), both of which are well-established amyloid-binding dyes ^29–32^, specifically bind the pharyngeal cuticle ^9^. Similarly, the blue-fluorescent Calcofluor white (CFW) and the yellow-red-fluorescent Eosin Y (EY), which bind chitin ^33^ and its deacetylated derivative chitosan ^34^ respectively, specifically bind the pharyngeal cuticle ^9^.

We incubated worms in wact-190 and wact-390 (a second crystalizing molecule) for 3 hours followed by 2 hours of incubation of the dyes. We found that the amyloid dye patterns are often disrupted by the nascent objects (Figure 4). We also observe a similar pattern in the background where a single IDR-rich protein (ABU-14) is tagged with GFP ^10^ (Supplemental Figure 3). By contrast, the chitin and chitosan dyes are relatively unperturbed (Figure 4; Supplemental Figure 3). In some instances, the birefringent crystals overlap with amyloid stain (blue arrows, Figure 4). In other cases, there is little overlap between the birefringent signal and the amyloid stain (orange arrows, Figure 4).

**Figure 4.**
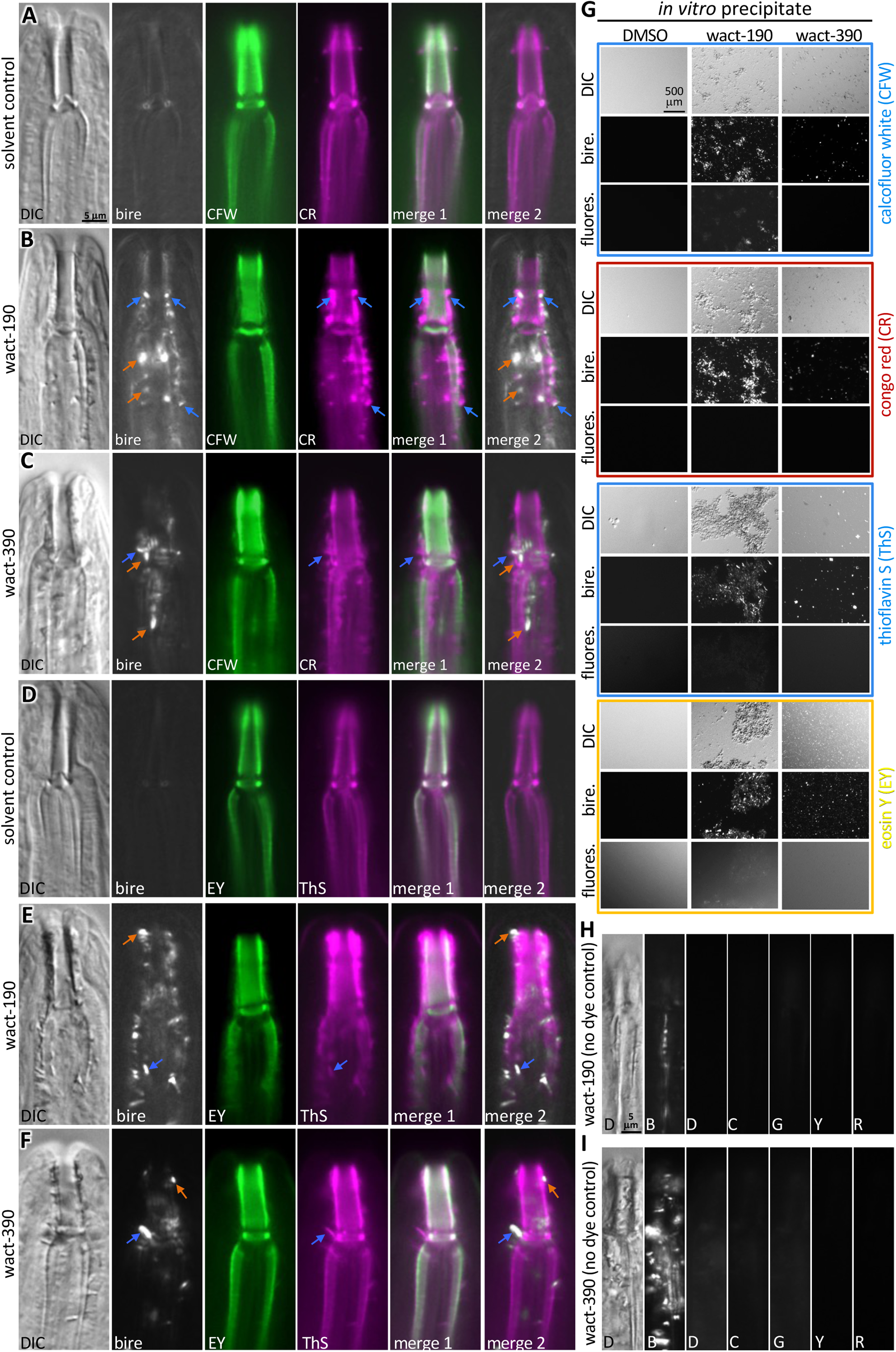
Crystals Preferentially Disrupt Amyloid Dye-Staining Material of the Pharyngeal Cuticle. **A-F**. Images of the anterior-most end of the worm. The imaging channel is indicated (lower left) as is the wactive treatment (left-hand side of micrograph series). DIC, differential interference contrast; bire, birefringence; CFW, calcofluor white; ThS, thioflavin S; EY, eosin Y. Merge 1 is the merge of the two dye signals. Merge 2 is the merge of the signals from the amyloid-binding dye and birefringence. Blue arrows indicate overlapping signals of the amyloid-binding dye and birefringence; orange arrows show a lack of amyloid-binding dye signal associated with birefringence signal. The scale in A applies to A-F. **G.** Control for the ability of dyes to bind wactive crystals. Precipitate of wact-190 or wact-390 was formed in solution (or DMSO solvent control) and was subsequently stained with the dye indicated on the right of each set. The dyes were used at the same concentration as the *in vivo* staining experiments as described in the methods. Each condition was imaged using DIC, birefringence (bire.) or the appropriate fluorescent (floures.) channel, as indicated on the left of each panel. The dyes were not rinsed from the slide, which accounts for the non-specific signal in the EY fluorescent channel. The scale in the upper left micrograph applies to all images in G. **H-I.** Controls for the fluorescence of crystals in the different channels. D, differential interference contrast (DIC); B, birefringence; C, CFP; G, GFP; Y, YFP; R, RFP channels. The scale bar in H also applies to I. 60 μM of wact-190 and wact-390 was used in all experiments relevant to panels A-I.

To control for the possibility that the dyes bind the crystals, we asked whether the dyes bind to *in vitro* precipitated small molecules. None of the dyes fluoresced brightly in association with the *in vitro* precipitated small molecules (Figure 4G). We also examined the dye-staining patterns in the presence of much larger 24-hour old *in vivo* crystals. We found many obvious examples where the larger birefringent signals do not overlap with the CR signal (orange arrows, Supplemental Figure 4). These results suggest that nascent crystals can displace the CR-stained material, which often appears to surround the small crystals. This is consistent with the matrix-like material that surrounds the objects in the TEM images (see Figure 2h for example in ^6^. As the crystals grow, their larger mass becomes more distinct from the cuticle and their intimate association with the amyloid-staining material is often lost.

### Amyloids Likely Seed the Formation of Small Molecule Crystals

As shown above, crystal-forming small molecules are rich in amyloid-binding substructures and that crystals can perturb the amyloid-like material in the pharynx cuticle as they grow. These observations raised the possibility that the amyloid-like material in the pharynx cuticle ^9^ might seed the formation of the small molecule crystals. If true, then coating the amyloid-like material with well-characterized amyloid-binding dyes might block crystal formation. We tested this idea by first asking whether wact-190-associated lethality of *C. elegans* is suppressed by co-incubating the worms with the amyloid-binding dyes CR and ThS in our standard 6-day viability assay ^5^. Indeed, we found that CR and ThS robustly suppress wact-190 lethality whereas the chitin/chitosan-binding dyes do not (Figure 5A). Second, we asked whether the canonical amyloid-binding dyes can suppress wact-190 crystal formation. We incubated CR and ThS, along with CFW and EY with the animals either for 3 hours before incubating the worms with wact-190 for another 3 hours (i.e. a pre-incubation) or at the same time as a 3-hour incubation with wact-190 (i.e. a co-incubation). We found that both CR and ThS suppress crystal formation in both incubation schemes, but that the CFW and EY dyes do not (Figure 5B and 5C). These data suggest that the crystals formed from the exogenous compounds may be seeded by the amyloid-like material in the pharynx cuticle, which can be blocked by other amyloid-interacting compounds.

**Figure 5.**
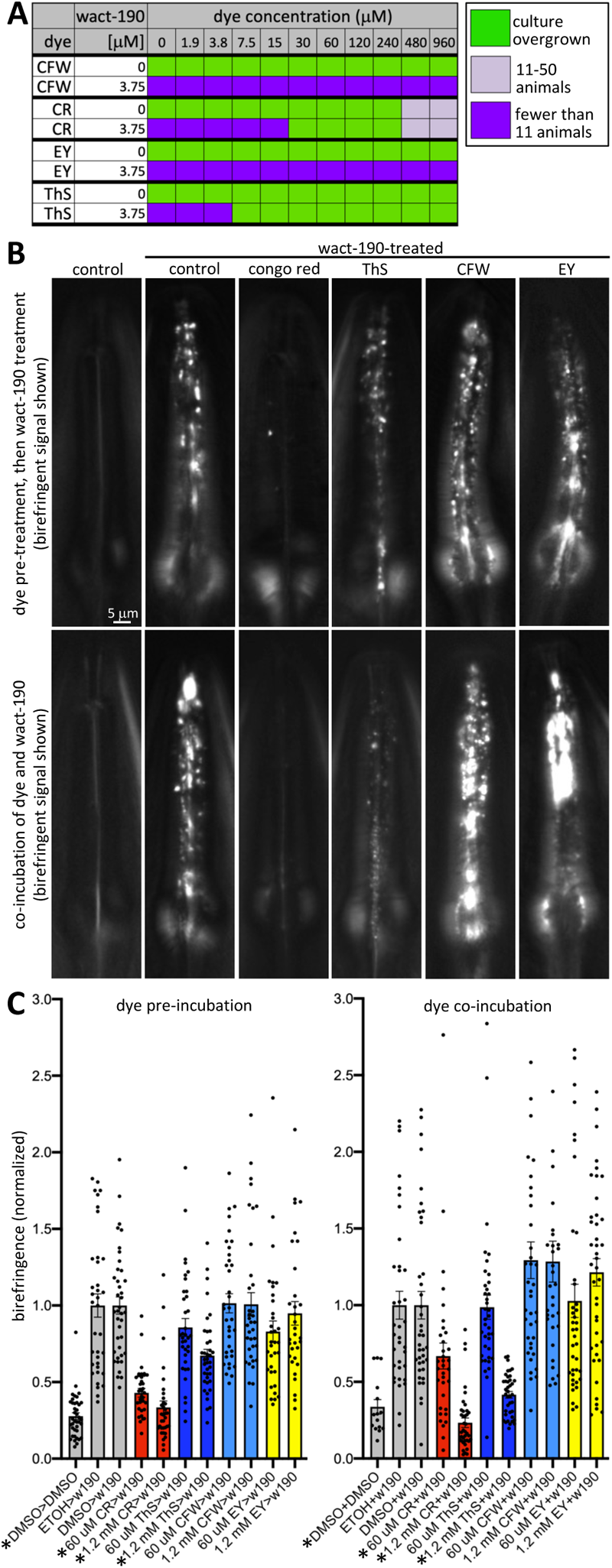
Amyloid-Binding Dyes Suppress Crystal Formation. **A.** Six-day viability assay results illustrating that amyloid binding dyes (CR and ThS) can suppress wact-190-induced lethality, but that chitin/chitosan-binding dyes (CFW and EY) cannot. Each cell shows the summary of 3 trials each done in quadruplicate. **B-C.** Amyloid-binding dyes, but not chitin/chitosan-binding dyes, can suppress crystal formation. On the top row, animals were pre-incubated in the indicated dye or solvent control for 3 hours. Then, the right-most five samples were incubated in 60 μM wact-190 for three hours without the initial dyes present. On the bottom row of (B), animals were incubated in the indicated dye or solvent control, and for the five right-most five samples, simultaneously incubated with wact-190 (60 μM) for 3 hours. Representative birefringence images are shown in (B) and the corresponding signal quantification is reported in (C). The scale bar in the upper left of B applies to all images in the series. Samples labelled with an asterisk indicates a significant difference relative to control (*p*<0.01). Standard error of the mean is shown.

We further investigated the idea that amyloids might seed the small molecule crystals by asking whether Aβ42 fibrils could seed the formation of wact-190 objects *in vitro*. We prepared solutions of Aβ42 at 25 μM and wact-190 at 25, 0.5 and 0.1 μM in PBS and incubated them overnight either alone or in combination with each other at different ratios. The next day, we examined the self-assembled structures by TEM (see Methods). We found objects associated with the Aβ42 fibrils were present in solutions with wact-190 present in either 10-fold more (Figure 6D), 2-fold more (Figure 6C), or 50-fold less than the concentration of Aβ42 (Figures 6G-H), but not 250-fold less (Figures 6K-L) or without wact-190 present (Figures 6A, E, and I). We infer that the objects are self-assemblies of wact-190. Together with the *in vivo* data presented above, these data suggest that amyloids are capable of seeding wact-190 self-assembly into higher order structures.

**Figure 6.**
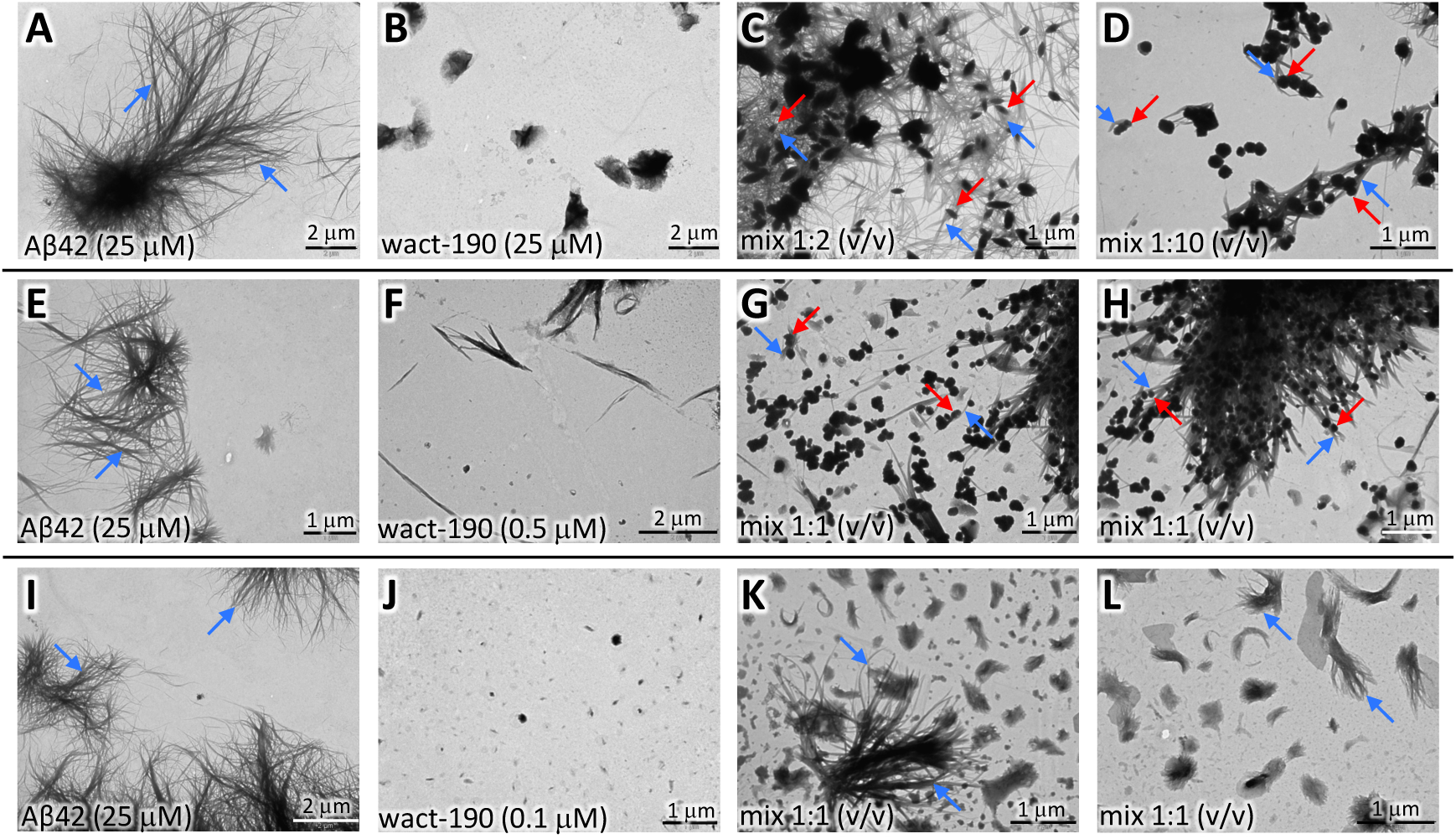
Aβ42 Fibrils can Seed wact-190 Objects. Transmission electron microscopy (TEM) images of solutions allowed to self-assemble overnight. **A-D.** Mixing solutions of 25 μM Aβ42 and 25 μM wact-190 in PBS. Unmixed 25 μM Aβ42 (A) and 25 μM wact-190 (B); 1-fold Aβ42 diluted with 2-fold wact-190 (to a final concentration of 8.33 μM Aβ42 and 16.66 μM wact-190) (C); 1-fold Aβ42 and 10-fold wact-190 (to a final concentration of 2.27 μM Aβ42 and 22.72 μM wact-190) (D). **E-H.** Mixing solutions of 25 μM Aβ42 and 0.5 μM wact-190 in PBS. Unmixed 25 μM Aβ42 (E) and 0.5 μM wact-190 (F); 1-fold Aβ42 diluted with 1-fold wact-190 (to a final concentration of 12.5 μM Aβ42 and 0.25 μM wact-190)(G and H). **I-L.** Mixing solutions of 25 μM Aβ42 and 0.1 μM wact-190 in PBS. Unmixed 25 μM Aβ-42 (I) and 0.1 μM wact-190 (J); 1-fold Aβ42 diluted with 1-fold wact-190 (to a final concentration of 12.5 μM Aβ42 and 0.05 μM wact-190) (K and L). Examples of Aβ42 amyloid fibrils are indicated with a blue arrow and examples of presumptive wact-190 objects that are associated with Aβ42 fibrils are indicated with a red arrow.

### Amyloid Disruptors are Revealed in a Screen for Small Molecule Suppressors of Crystal Formation

Above, we show that amyloid-binding probes can block crystal formation. This raised the possibility that screens for small molecules that disrupt crystal formation might yield new amyloid-interacting and/or disrupting molecules. To investigate, we screened Microsource’s Spectrum library of 2560 drugs and natural products for those that might suppress crystal formation.

We screened the Spectrum library in our standard six-day liquid viability assay whereby multi-well plates are seeded with first larval stage animals (see methods and ^2^). We previously established that wact-190’s lethality in young larvae is dependent on crystal formation ^5^. Control (solvent-only) wells are overgrown with worms after six days (Supplemental Table 5). By contrast, wells with wact-190 (3.75 μM) contain either arrested or dead L1s at the assay endpoint (Supplemental Table 5 and ^5^).

Screening the Spectrum library molecules at 60 μM yielded 85 hits that: i) suppress wact-190-induced lethality; ii) suppress wact-190 crystal formation (*p*<0.05); and iii) fail to suppress lethality induced by a counter-screen control that does not form crystals (the succinate dehydrogenase inhibitor wact-11 ^2^) (Supplemental Table 5). To assess the structural diversity of the suppressors, we built a structural similarity network of the molecules (Figure 7A). A Tanimoto coefficient cut-off of 0.8 and manual inspection revealed 45 distinct core scaffolds, 17 of which are represented by multiple molecules (Figure 7A-B).

**Figure 7.**
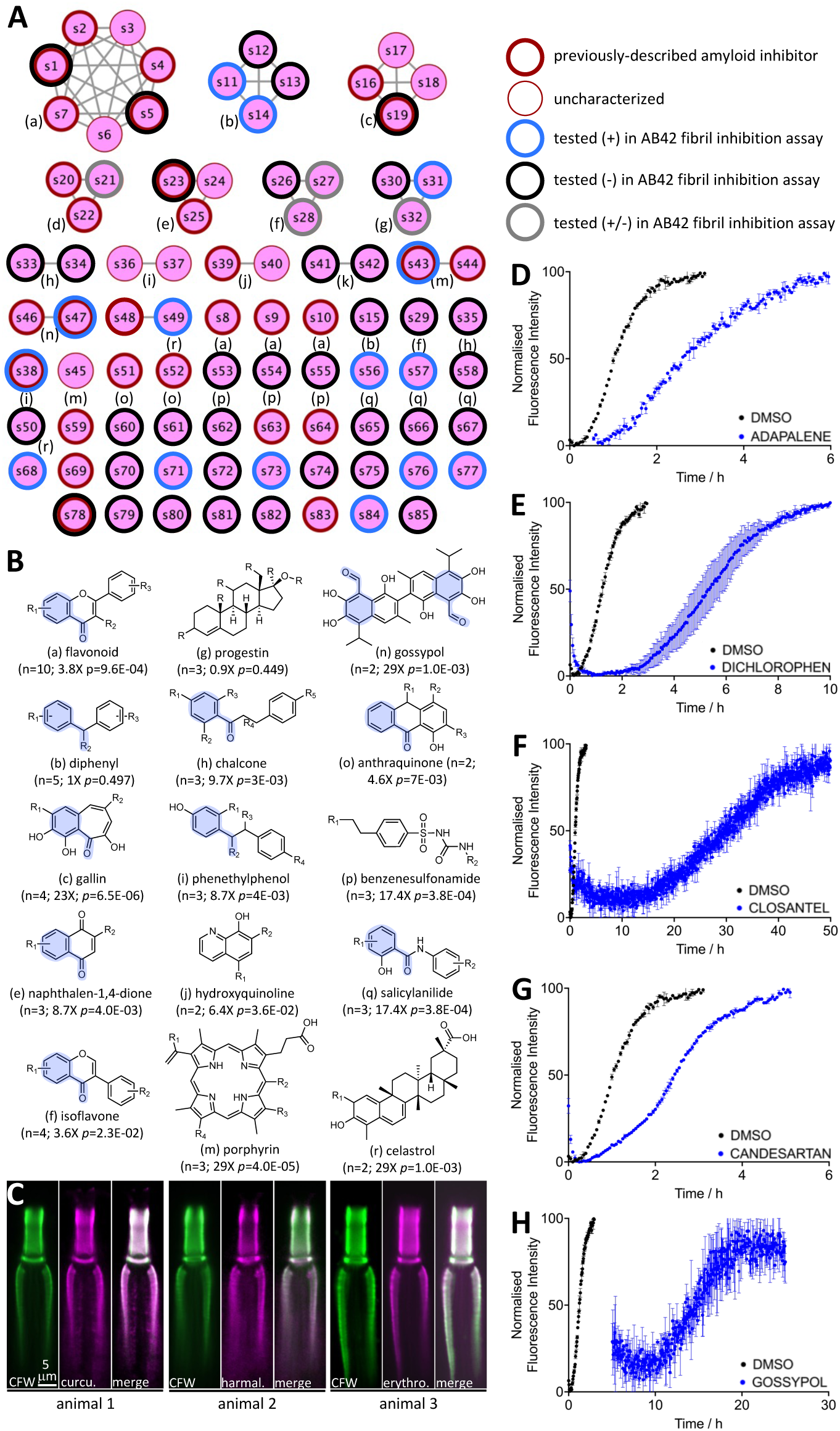
A Screen for wact-190 Suppressors Reveals Amyloid-Disrupting Molecules. **A.** A structural similarity network of the 85 Spectrum library hits that suppress wact-190 lethality and crystal formation. The nodes (circles) represent a drug or natural product. The links (straight lines) represent a Tanimoto score of structural similarity of 0.8 or more (see Methods). A bold red outline indicates known disruptors of amyloid fibril formation. See Supplementary Table 3 for details and names of the compounds. **B.** Core scaffolds among the drug suppressors of crystal formation. The small case letter preceding the scaffold name corresponds to the nodes indicated in (A). The number (n) of individual molecules within the network is indicated, as is the enrichment of that scaffold within the network relative to that expected by random chance based on the prevalence of that scaffold within the entire library (the hypergeometric *p* value is shown). The benzaldehyde substructure is highlighted in blue. **C.** Examples of the localization of crystal suppressors (60 μM) that are fluorescent. Calcofluor white (CFW) is used as a counter-stain control; curcu., curcumin; harmal., harmaline; erythro., erythrosine. See Supplemental Figure 5 for additional examples and controls. The scale in panel C applies to all images in the series. **D-H.** *In vitro* assays to characterize the ability of the indicated molecule to inhibit Aβ42 fibril formation in secondary nucleation assays (see methods, Supplementary Table 5 and Supplementary Figure 8 for additional details). Adapalene is a known standard inhibitor of Aβ42 fibril formation ^61^.

Informatic analyses of the 85 suppressors revealed that 15 of the 45 scaffold groups (33%) contained at least one molecule known to play a direct role in disrupting amyloid fibril formation (bold outline in Figure 7A; Supplemental Table 5). Most of these scaffolds are enriched among the suppressors relative to the entire Spectrum library (see Figure 7B for *p* values). Of note, a benzaldehyde substructure is significantly enriched among the suppressors (2.8-fold enrichment, *p*=2.3E-14; blue highlight in Figure 7B). Importantly, we found that several of the suppressors are fluorescent and of these, all specifically stain the pharyngeal cuticle (Figure 7C and Supplemental Figure 5).

We investigated whether the set of 85 wact-190 suppressors is enriched for amyloid disruptors relative to the molecules within the Spectrum library. The lack of a comprehensive database of previously characterized small molecule amyloid disruptors precluded a systematic analysis. We therefore randomly selected 100 molecules from the remaining Spectrum molecules to further investigate. We used the same analytical approach that we employed for the 85 Spectrum suppressors. We first assembled the random molecules into a structural similarity network, revealing 75 structurally-distinct scaffolds (11 clusters plus 64 singletons) (Supplemental Figure 6A). We then investigated the literature for evidence of the ability to disrupt amyloid fibril formation for each of the 100 random molecules (see methods). Two of the 75 core scaffolds (3%) had one molecule shown to be a disruptor of amyloid fibril formation (Supplemental Figure 6B). From this limited analysis, we found that the 85 Spectrum suppressors are over 10-fold enriched in amyloid disruptors relative to the randomly selected set.

### Small Molecule Suppressors of Crystal Formation Inhibit Aβ42 Fibril Formation

The above results suggest that the mechanism by which molecules suppress crystal formation is intimately related to amyloid disruption. We therefore asked whether any of the molecules for which we could not find literature-based evidence for amyloid disruption have such capability. While different suppressors have been shown to disrupt different types of amyloids (Supplemental Table 5), we focused on Aβ42 for reasons of convenience ^35,36^. We measured the incubation time by which Aβ42 fibril formation reaches saturation and asked which molecules are capable of significantly (*p*<0.05) extending the half-time of saturation by at least 50% in a standard assay ^35^. Of the eight molecules previously shown to disrupt amyloids, 3 inhibited Aβ42 fibril formation in our assay (Figure 7A, Supplemental Table 5), indicating that not all published results translate to this specific assay. Of the 44 unknowns tested, 11 (25%) inhibited Aβ42 fibril formation (Figures 7A, D-H, Supplemental Table 5). Follow up analyses on the hits shown in Figure 7E-H shows that Candesartan is a specific nucleation inhibitor while the other molecules inhibit either nucleation and/or elongation. Importantly, all of these inhibitory mechanisms require fibril binding (Supplemental Figure 7). Hence, the ability of a small molecule to suppress crystal formation *in vivo* is predictive of its ability to inhibit amyloid fibril growth *in vitro*.

### A Scaffold’s Ability to Crystalize or Suppress Crystallization is Interchangeable

A comparison of the structures of crystallizers and suppressors revealed that the former have fewer hydrogen-bond donors than the latter and completely lacked hydroxyl groups (*p*<1E-10) (Figure 8A-B). The crystallizers are also more halogenated than the suppressors (Figure 8C). We also found that some crystallizers share the identical scaffold as some suppressors. For example, the scaffolds of the wact-128 and wact-498 crystallizers are identical to that of oxyclozanide and koparin crystal suppressors, respectively (Figure 8D). In addition, the scaffold of the wact-415 crystallizer is similar to that of ThS, which also suppresses crystal formation (Figure 8D). These observations raised the possibility that both classes of molecules (crystallizers and suppressors) compete for the same environment within the cuticle.

**Figure 8.**
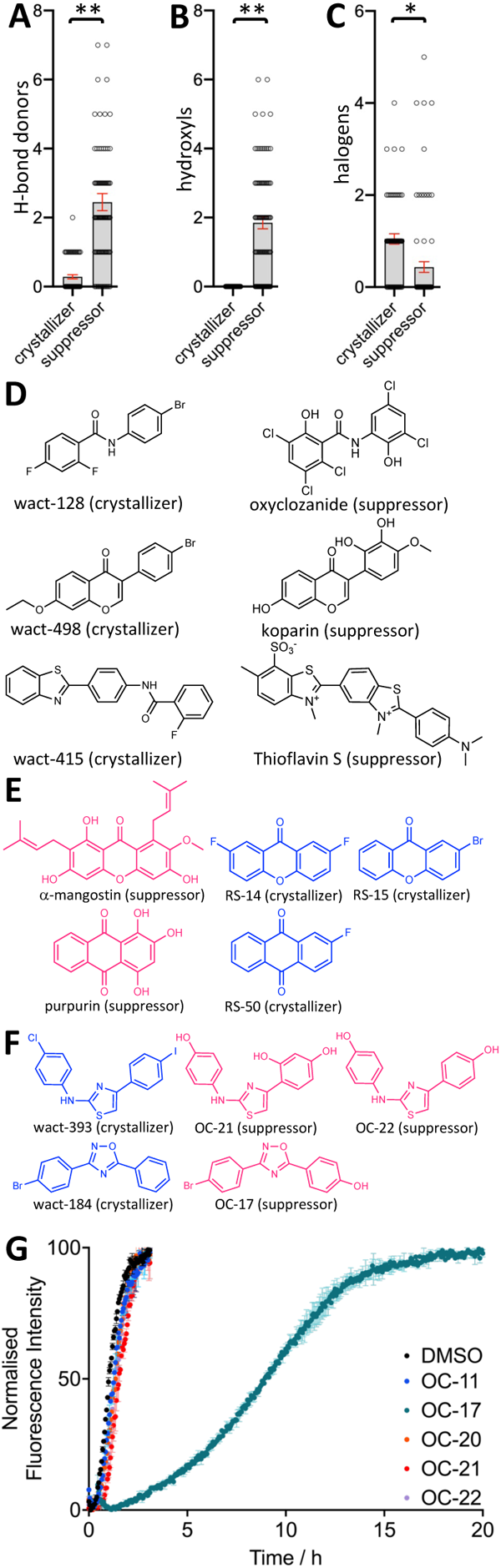
Crystallizers and Suppressors can Share Identical Core Structures. **A-C**. Physicochemical features of crystallizers and suppressors. A single and double asterisk indicate *p*<0.001 and *p*<1.0E-10, respectively. The standard error of the mean is shown. **D.** Crystallizers and suppressors sharing an identical or similar core structure. **E.** Core scaffolds of suppressing molecules that can be turned into crystallizers (see Supplemental Table 6 for additional details). **F.** Core scaffolds of crystallizing molecules that can be turned into suppressors (see Supplemental Table 7 for additional details). **G.** *In vitro* assays to characterize the ability of the indicated suppressor molecule to inhibit Aβ42 fibril formation in secondary nucleation assays (see methods and Supplementary Figure 8 for additional details).

We tested whether we could convert a crystalizing molecule into a suppressor, and a suppressor into a crystallizer, by altering the abundance of hydroxyl and halogen decorations on the respective core structures. To do this, we searched the ZINC database of the world’s commercially available molecules ^37^ for close structural analogs of the 85 suppressors that lacked hydroxyl groups and are halogenated. We screened 33 available analogs of 6 different suppressors. We found that 11 (33%) of these molecules induced object formation within the pharynx in at least 25% of the population (Supplemental Table 6; Figure 8E). This percentage is comparable to the 36% of molecules that induce object formation from our heavily biased collection of worm bioactive small molecules ^5^. Second, we tested 25 hydroxylated analogs of 10 different crystalizing small molecules for their ability to suppress the lethality associated with crystal-forming wact-190. Five of the 25 hydroxylated crystallizers (20%) suppress wact-190 lethality, and all of these suppress wact-190 crystal formation (*p*<0.05) (Supplemental Table 7; Figure 8F). We conclude that select core small molecule structures can be pushed towards crystal formation or its suppression by changing the relative abundance of halogens and hydroxyls that decorate the molecule. These data further supports the model that crystallizers and suppressors likely compete for the same niche.

Finally, we tested the five converted suppressor molecules in the Aβ42 fibril formation assay and found that the most robust suppressor (OC-17) robustly inhibited Aβ42 fibril formation (Figure 8G; Supplemental Table 7; Supplemental Figure 7E). Hence, converting crystallizers into suppressors is another route with which to identify amyloid disruptors.

## Discussion

### Insights into Crystal Formation within the *C. elegans* Pharynx Cuticle

Here, we have described our investigation into why some small molecules crystalize within the cuticle of the *C. elegans* pharynx. We found that: i) many crystal-forming molecules have a core scaffold known to bind amyloid; ii) crystals can displace amyloid-like material within the pharyngeal cuticle; iii) other molecules that are known to bind amyloids can block crystal formation; iv) molecules that block crystal formation specifically localize to the tissue where crystals form; and v) Aβ42 fibrils can seed the formation of wact-190 objects *in vitro*. Together with our previous work that shows the pharynx cuticle to be rich in amyloid-like material ^9^, these observations suggest that the amyloid-like material within the pharynx cuticle seed the formation of crystals.

We can speculate in general terms as to how the amyloid-like material in the pharynx cuticle might seed small molecule crystals. Our physicochemical analyses of crystalizing compounds show that they are rich with π bonding capability, hydrogen bond acceptors, and polar surface areas (this work and ^5^). Indeed, our *in vitro* analysis also shows that π bond interactions likely play a key role in the ability of these molecules to crystalize within the pharynx. Concomitant with its amyloid-like nature ^38,39^, pharynx cuticle proteins are highly enriched in π-bonding capability relative to other tissue proteomes in *C. elegans* and is only surpassed by the collagens ^9^. Furthermore, the pharynx secretome is uniquely rich with polar uncharged residues with hydrogen bond donor capability (i.e., Ns, Qs, Ss, Ts, and Ys) ((*p*<1E-11) ^9^). Hence, the unique complementarity between the proteins within the pharyngeal cuticle and the crystal-forming compounds likely facilitates the seeding of small molecule crystals.

Small molecule crystallization is restricted to the pharyngeal cuticle ^5^, yet the collagens (which are highly abundant within the body cuticle) are also rich in π-bonding capability ^9^. Hence, there are likely additional properties of the pharynx cuticle beyond its amyloid-like nature that creates the unique niche that facilitates crystal formation. Indeed, our recent work shows that polar lipids within the pharynx cuticle likely act as a sink to concentrate the hydrophobic molecules that go on to crystallize ^6^.

A model of crystal formation emerges upon considering these and other observations. First, polar lipids within the pharynx cuticle likely act as a sink to concentrate hydrophobic small molecules, including those that go on to crystallize ^6^. Once concentrated, the amyloid-like material within the pharynx cuticle seeds the crystallization of small molecules that are capable of interacting with the amyloid-like material. Many of the molecules that suppress crystal formation have similar properties to the crystallizing molecules except that they are rich in hydroxyl groups and therefore cannot crystalize themselves. Instead, they mask the niche to prevent the seeding of crystals from small molecules that would otherwise do so.

### Crystal Formation can be Exploited to Identify Candidate Amyloid Inhibitors

Through this investigation, we discovered that the phenomenon of small molecule crystal formation can be readily exploited to identify candidate inhibitors of amyloid growth. Our screen of just over 2500 small molecules for those that can suppress crystal formation yielded 85 reproducible hits. One third of these were previously known inhibitors of amyloid formation. For example, the heme and chlorophyl precursor protoporphyrin IX, its synthetic derivative haematoporphyrin, and chlorophyllide-Cu are the only porphyrins contained within the Spectrum library, and all three were found to suppress crystal formation. Multiple studies have shown that porphyrins suppress amyloid formation both *in vivo* and *in vitro* and act through direct interaction with amyloidogenic proteins ^40–42^. The porphyrin ring is thought to disrupt the intrapeptide π-π interactions of the amyloidogenic peptide that is necessary to achieve superstructure ^42^. The porphyrins may suppress crystal formation by disrupting the ability of the amyloid-like material to seed crystals or may disrupt the small molecule crystallization itself.

A second common suppressor class are the flavonoids, which are a large class of polyphenolic secondary metabolites synthesized by plants ^43^. Many flavonoid structures have been shown to antagonize amyloid formation *in vitro* ^44–46^, in culture ^47^, and *in vivo* ^48–50^. Poor bioavailability may contribute to a lack of flavonoid-based therapies in the clinic, although converting anti-amyloid flavonoids into prodrugs may circumvent absorption barriers ^51^. Flavonoid interaction with Aβ42 is driven by hydrophobic and hydrogen bond interactions that antagonize the ability of Aβ42 to adopt a β-sheet secondary structure ^52^. Whether the flavonoids interact with amyloidogenic proteins within the cuticle in a similar way to suppress crystallization remains to be determined.

Upon testing 44 small molecules not known to us at the time to disrupt amyloid formation, we found that 25% disrupt Aβ42 aggregation. This hit rate is far higher than that previously reported for Aβ42 (0.4%) ^16^ or α-synuclein (<0.01%) ^18^. Notably, *C. elegans* screens for suppressors of crystal-induced death are inexpensive and high-throughput. Furthermore, the *in vivo* environment in which suppressors must act is complex and, relative to other *in vitro* assays, may better approximate the environment in which anti-amyloid compounds must act. Hence, screens for molecules that suppress crystal-induced lethality in *C. elegans* represent a novel and efficient approach to enrich for molecules that might have utility against human amyloid formation.

## Methods

### *C. elegans* Culture, Strains and Microscopy

*C. elegans* strains were cultured and synchronized as previously described ^5^. Six day viability assays were conducted as previously described ^2^. The wild type N2 Bristol strain was used (obtained from the *C. elegans* genetic Centre); NQ824 qnEx443[Pabu-14:abu-14:sfGFP; rol-6(d); unc-119(+)], which was a kind gift from David Raizen ^10^, was integrated into the genome in our lab as *trIs113*. For all imaging, a Leica DMRA compound microscope with a Qimaging Retiga 1300 monochrome camera was used. Before mounting worms on a 3% agarose pad for imaging, worms are washed 3X with M9 buffer and paralyzed with 50 mM levamisole or 30 mM NaN_3_. Dye staining experiments were performed as previously described ^9^.

### Identification of Crystallizing Small Molecules and Crystal Counts

The RLL1200 library of 1178 commercially available analogs of *C. elegans*-lethal molecules was assembled and purchased from ChemBridge (Supplemental Data File 1). This library was screened at 30 µM in liquid NGM media against synchronized *C. elegans* L1 larvae and *Rhabditophanes diutinus* ^53^ as previously described ^2^, with approximately 20 L1s plated per well of a 96-well plate. The final volume in each well was 100 µL and the DMSO concentration was 0.3% v/v. After 6 days of incubation at 25°C the number of living worms in each well was quantified. Molecules that killed *C. elegans* N2 and *Rhabditophanes* were tested for resistance in *C. elegans sms-5(ok2498)* mutants at 1.9, 7.5, and 30 µM. Crystal counts were performed as previously described ^5^.

### Characterization of the Structure of wact-190 and wact-416 *in vitro* Crystals

Both wact-190 and wact-416 crystal powder samples were deposited on a quartz zero-background sample holder. The sample was loaded in a 0.7 mm diameter capillary, to avoid preferment orientation. The powder X-ray diffraction (PXRD) pattern was collected using a Bruker D8 Discover diffractometer (Bruker, Germany) equipped with Goebel’s mirrors to create a θ:θ parallel beam geometry with a sealed X-ray tube source with a copper anode (40 Kv, 40 mA) and a LYNXEYE XE linear detector. The diffraction patterns were collected between 5 and 60° 2θ with step 0.02° 2θ for 4 seconds per step^54^. The cell indexing was performed by TOPAS 5.0 software, using the LSI-Index algorithm ^55^. The structure resolution was obtained by using the simulated annealing method in EXPO2014 ^56^. The founded structure was refined by the Rietveld Method using GSAS-IIin order to precisely refine the atomic positions ^57^. The CIF files obtained from the above method and structures were solved using Mercury software.

### Transmission Electron Microscopy (TEM) of Aβ42 and wact-190 *in vitro* Crystals

5 mM of each of Aβ42 and wact-190 were dissolved in pure DMSO and heated at 55 °C for 2 minutes for complete dissolution. Thereafter, all molecules were further diluted in 1X PBS to prepare a final concentration of 25 μM. This solution mixture was further heated at 90 °C for 3 hours, followed by a vortex and the mixture was then allowed to gradually cool down overnight that resulted in self-assembled structures in some cases. The combinations used for Aβ42 plus wact-190 were 1:1, 1:2, 1:5, and 1:10 ratios. Uncombined Aβ42 and wact-190 were used as controls (25 μM in 1X PBS). The self-assembled samples of all three (5 μL, 25 μM in 1X PBS) were drop cast on a 400-mesh carbon-stabilized Formvar-coated Cu grid (Ted Pella, California, USA). The sample was allowed to bind the surface for 2 minutes and then the excess sample was removed using a lint-free tissue, dried at room temperature, and imaged. Sample morphology was visualized using a JEM-1400 TEM (JEOL, Tokyo, Japan) accelerating at 80 kV ^54^.

### Identification of Chemical Suppressors of Crystal-Induced Death and Crystal Formation

To identify chemical suppressors of crystal-induced death, the Spectrum Collection (MicroSource Discovery Systems Inc.) of 2560 approved drugs and natural products was screened at a concentration of 60 µM in the background of 3.75 µM wact-190. This concentration of wact-190 results in crystal formation in the pharynx causing *C. elegans* N2 to arrest at the L1 stage. This screen was performed in liquid NGM media as previously described ^2^, with approximately 20 N2 L1s plated per well of a 96-well plate. The final volume in each well was 50 µL and the DMSO concentration was 0.8% v/v. After 6 days of incubation at 20°C the number adult and larval worms that have progressed from arrested L1 stage was quantified. Hits that suppressed the wact-190 crystal-induced L1 arrest were retested against solvent control, wact-190, and the non-crystal forming lethal molecule wact-11 at 10 µM (see ^2^) using the same method described above. Hits were further investigated for their ability to suppress crystal formation by co-incubating 3.75 mM of wact-190 and 60 mM of the suppressor with wild type animals in 96-well plate format. The fraction of animals with birefringent crystals was then counted at 100X magnification 48 hours later, counting 20 animals per sample per trial on average.

### Chemoinformatic Network Analysis, Network Display, Small Molecule Art, and Informatic Analyses

Structural similarity networks were constructed based on pairwise similarity scores calculated as the Tanimoto coefficient of shared FP2 fingerprints. Pairwise similarity scores were calculated using OpenBabel (http://openbabel.org) and the networks were visualized using Cytoscape ^58^ as previously described ^2^. For each network generated, a range of Tanimoto coefficient cut-offs were tested to determine which cut-off yielded a network that clustered obvious structural analogs together as determined by manual inspection. Suppressors were analysed for the presence of substructures (Figure 4b) using Filter-it version 1.0.2 from Silicos-it. Structures were drawn using ChemDraw (PerkinElmer Informatics). A random Spectrum network (Supplemental Figure 5) was generated by assigning all molecules within the Spectrum library a random number using excel, then sorting the molecules in order of the random number, then choosing the top 100 molecules that did not include any of the 85 Spectrum suppressors.

We investigated whether the crystalizing molecules and a randomly selected set of true negatives (that fail to robustly form crystals ^5^) were previously characterized to interact with amyloids using the SciFinder Scholar tool ^21^. To do this, we first assembled the crystalizing molecules into 29 groups based on distinct scaffolds and ensured that our set of 62 true negatives represented 62 distinct core scaffolds (within the set of true negatives). We then reduced each group (or single molecule if it was not part of a group) to its core scaffold structure by removing hydroxyl groups, halogens, or substructures that were not in common to all members of the group. With the resulting core scaffold structures, we first searched SciFinder for any molecule that contained that core scaffold. With these resulting molecules, we then searched the associated biological literature abstracts within SciFinder for the term ‘amyloid’. Because of the simplicity of some core scaffolds, the number of respective analogs and associated abstracts numbered in the tens of thousands. In these cases, we instead first chose one molecule from the group (or the single molecule if it was not part of a group) and searched SciFinder for analogs with >80% identity. This reduced the number of relevant structures and associated literature to search and kept the focus on closely related structures. Which of the two tactics was employed is indicated in the relevant Supplementary Table and Supplementary Data File for each scaffold group (or single molecule). Abstracts containing the term ‘amyloid’ were read (and where unclear, the associated literature was read) to ensure that evidence of either direct amyloid interaction *in vitro* was being measured or that if *in vivo* samples were being analyzed, staining that was coincident with amyloid structures was being measured. Evidence relying solely on the reduction of amyloids *in vivo* were not considered ‘hits’ because of potential indirect mechanisms. The PubMed IDentification numbers (PMIDs) and/or patent numbers of the ‘hits’ are reported in the associated Supplementary Tables and Supplementary Data File.

We analyzed both the list of 85 Spectrum suppressors of crystal formation and the 100 randomly chosen molecules by searching that compound and the word ‘amyloid’ in both a PubMed search and a Google search. We inspected the search results for evidence that the molecule of interest had a demonstratable inhibitory effect on amyloid formation *in vitro*, with or without supporting *in vivo* evidence. *In vivo* or *in silico* evidence of a decrease in fibril abundance without evidence of a direct effect was excluded from consideration. Molecules that lacked evidence for a direct effect, but had close structural analogs that have a demonstratable inhibitory effect on fibral formation were also excluded from further consideration.

To identify hydroxylated analogs of the crystallizing molecules and reduced analogs of the suppressors, Open Babel ^59^ was used for the analog searches. Using SmiLib v2.0 ^60^, a virtual library of approximately 55,000 variably hydroxylated analogs of 161 distinct crystallizing molecules was generated, as well as a virtual library of approximately 950,000 de-hydroxylated and variably halogenated analogs of 12 distinct crystal suppressor molecules. The 55K and 950K virtual libraries were de-duplicated and independently combined with de-duplicated and de-salted versions of MolPort’s All Stock Compounds Database and the ZINC In-Stock Database ^37^. A duplicate search of these combined datasets revealed 34 crystallizer analogs and 88 crystal suppressor analogs that were available for purchase from MolPort. These analogs were further filtered by price, real availability, redundancy among parent Spectrum suppressors, and aqueous solubility, leaving 26 crystallizer analogs and 33 crystal suppressor analogs to be assayed.

### Dye Suppression Assays

To test for the ability of dyes to suppress wact-190-induced worm lethality, 80 μL of NGM containing HB101 bacteria (OD_600_ = 1.8-2.2) and 3.75 µM wact-190 was added to each well of a 96-well plate, to which each dye was pinned (pinner model VP381N, V & P Scientific, Inc) to a final concentration of 0, 0.94, 1.88, 3.75, 7.5, 15, 30, 60, 120, 240, 480 and 960 µM, in quadruplicates. 20 µL of M9 containing synchronized wildtype L1s were then added to each well at ∼50 L1/well. Three sets of controls were simultaneously set up: wact-190 + DMSO (or 70% ethanol) (no dye), DMSO + dye (no wact-190) and DMSO + DMSO (or 70% ethanol) (no wact-190, no dye). All samples were set up in quadruplets (Day 0) and incubated at 20°C in a shaker at 200 rpm. Worm viability was scored on Day 6.

To test for the ability of dyes to suppress wact-190-induced crystal formation, synchronized wildtype young adults were added (∼ 1,000 worms in 50 µL) to a 1.5 mL Eppendorf tube containing 500 µL NGM buffer and bacteria. Adult worms were either (i) pre-incubated for 3 hours with 60 µM or 1200 µM of each dye or solvent (DMSO) only, rinsed 3X with M9 buffer then incubated with 60 µM wact-190 in a fresh Eppendorf tube, or (ii) co-incubated for 3 hours with 60 µM or 1200 µM of each dye (or solvent) and 60 µM wact-190. Crystal formation was recorded by observing the birefringence of the wact-190 crystals both in the presence and absence of the dye. wact-190 crystal formation was compared with a negative control (no wact-190), that yielded no such birefringence. All observations and imaging were carried out using 40X objective lens of Leica DMRA2 compound microscope. The amount of birefringence was quantified using Fiji/ImageJ software. 10-20 worms per sample were analyzed in 3 biological trials.

### *In vitro* Crystal Signal Emission Controls with and without Dye Staining

To test for signal emission of wact-190 or wact-390 crystals, as detected with our fluorescent microscopy filter sets, 3 µL of a 10 mM wactive stock was first added to 7 µL of water and mixed in a microcentrifuge tube, then directly added to a microscope glass slide and coverslip applied. The appearance and birefringence of the formed crystals were observed under differential interference contrast (DIC), birefringence, and fluorescence filter sets.

To test for whether the dyes specifically bind the *in vitro* precipitated crystals, the same procedure was followed as above, but each dye was added at a final concentration that was used in the dye staining experiments after the wactive precipitated in water. Correlation between crystal birefringence (bright white dots in a dark background) and dye fluorescence in their respective fluorescence channels was examined. All observations were made using the 10X objective lens of the Leica DMRA2 compound microscope.

### Aβ42 Aggregation Assay

#### Recombinant Aβ42 expression

The recombinant peptide MDAEFRHDSGY EVHHQKLVFF AEDVGSNKGA IIGLMVGGVV IA, here called Aβ42, was expressed in the *E. coli* BL21 Gold (DE3) strain (Stratagene, CA, U.S.A.) and purified as described previously ^35^. Briefly, the purification procedure involved sonication of *E. coli* cells, dissolution of inclusion bodies in 8 M urea, and ion exchange in batch mode on diethylaminoethyl cellulose resin followed by lyophilisation. The lyophilised fractions were further purified using Superdex 75 HR 26/60 column (GE Healthcare, Buckinghamshire, U.K.) and eluates were analysed using SDS-PAGE for the presence of the desired peptide product. The fractions containing the recombinant peptide were combined, frozen using liquid nitrogen, and lyophilised again.

#### Aβ42 aggregation kinetics and fibril preparation

Solutions of monomeric Aβ42 were prepared by dissolving the lyophilized Aβ42 peptide in 6 M guanidinium hydrocholoride (GuHCl). Monomeric forms were purified from potential oligomeric species and salt using a Superdex 75 10/300 GL column (GE Healthcare) at a flowrate of 0.5 mL/min, and were eluted in 20 mM sodium phosphate buffer, pH 8 supplemented with 200 µM EDTA and 0.02% NaN_3_. The centre of the peak was collected and the peptide concentration was determined from the absorbance of the integrated peak area using extinction coefficient ε280 = 1490 l Mol^-1^ cm^-1^. The obtained monomer was diluted with buffer to the desired concentration and supplemented with 20 μM Thioflavin T (ThT) from a 2 mM stock. Each sample was then pipetted into multiple wells of a 96-well half-area, low-binding, clear bottom and PEG-coated plate (Corning 3881), 80 µL per well, in the absence and the presence of different molar-equivalents of small molecules (1% DMSO). Assays were initiated by placing the 96-well plate at 37 °C under quiescent conditions in a plate reader (Fluostar Omega, Fluostar Optima or Fluostar Galaxy, BMGLabtech, Offenburg, Germany). The ThT fluorescence was measured through the bottom of the plate using a 440 nm excitation filter and a 480 nm emission filter.

### Statistics and Graphs

Except where indicated, statistical differences were measured using a two-tailed Students T-test. Dot plots were generated using Prism 8 graphing software.

### Animal Ethics Statement

We (the authors) affirm that we have complied with all relevant ethical regulations for animal testing and research. Given that our experiments focused exclusively on the invertebrate nematode worm *C. elegans*, no ethical approval was required for any of the presented work.

### Data Availability Statement

Any primary data generated or analyzed during this study that are not included in this published article and its supplementary information files are available upon request.

## Supporting information

Supplemental Datafile 1

## Acknowledgements

We thank David Hall, Zeynep Altun, and Chris Crocker at WormAtlas for permission to modify Worm Atlas schematics. WormAtlas is funded by NIH OD 01943 to D.H. Hall. This work is supported by CIHR grants (153024 and 173448) and a CRC to PJR, UKRI grants (10059436, 10061100) to MV and RIH, and a John Templeton Foundation Grant to DLB-Y and EG. The opinions expressed in this publication are those of the authors and do not necessarily reflect the views of the John Templeton Foundation.

## Author Contributions

PJR and MK found that crystals form in the pharyngeal cuticle. MK performed the microscopy and associated quantification. JK performed additional small molecule screens that revealed new crystal-forming compounds, performed the screen of the Spectrum library for suppressors of crystal-induced death and follow-up work, and performed the network analyses. ARB designed the novel 1,178-compound custom library and identified the crystallizer and suppressor analogs. RIH under the guidance and funding of MV, performed the Ab42 fibril inhibition assay. OST, under the guidance and funding of DLB-Y and EG, determined the structure of wact-190 and wact-416 *in vitro* crystals and showed that Aβ42 fibrils can nucleate wact-190 objects. DH determined whether the fluorescent Spectrum suppressors specifically stain the pharyngeal cuticle. PJR supervised, coordinated and secured funding for the project, composed the display items and wrote the manuscript with the help of co-authors.

## Competing Interests

RIH is a consultant of WaveBreak Therapeutics (formerly Wren Therapeutics). MV is a founder of WaveBreak Therapeutics.

---

**Supplemental Table 1.**
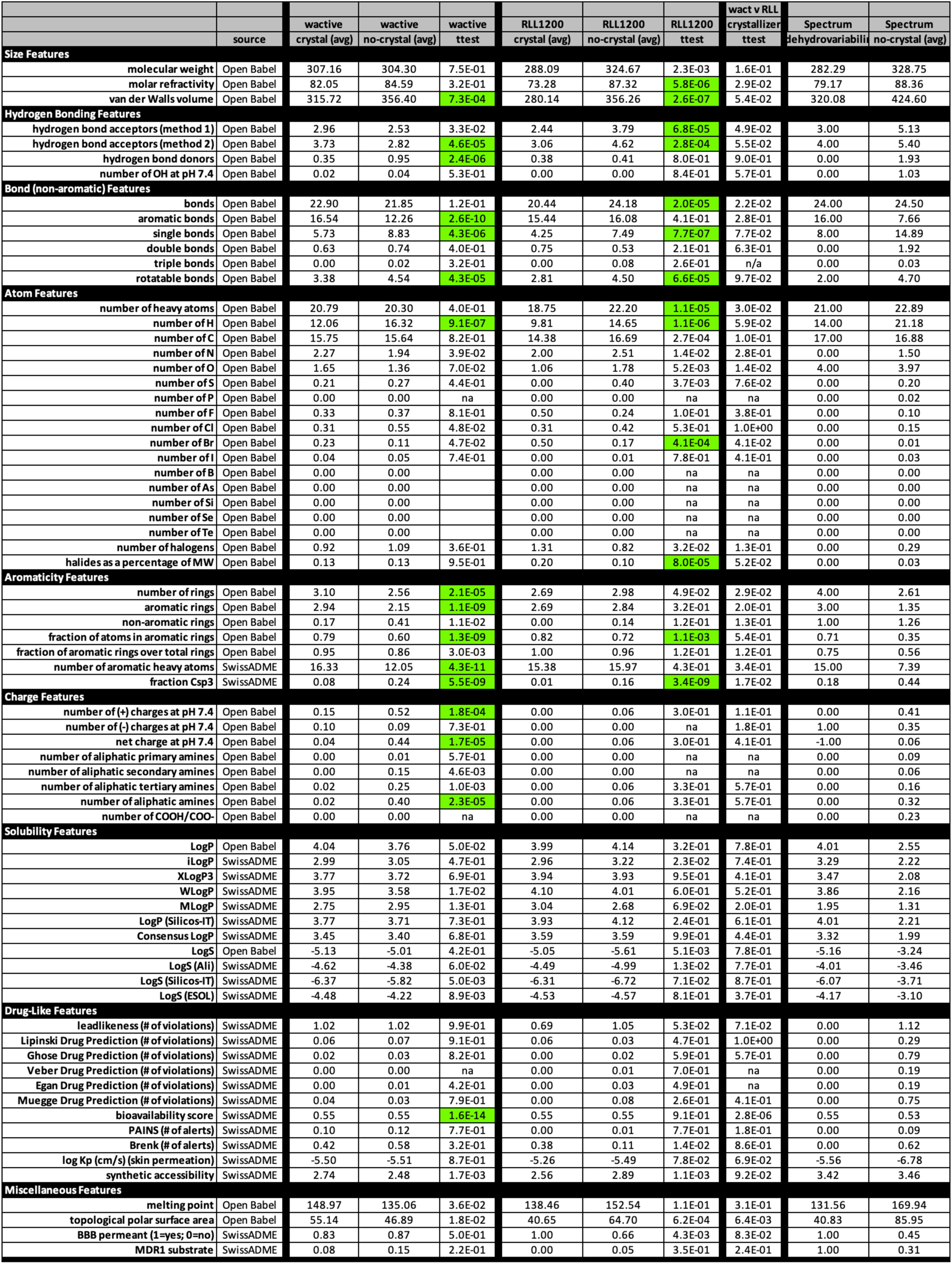

**Supplemental Table 2.**
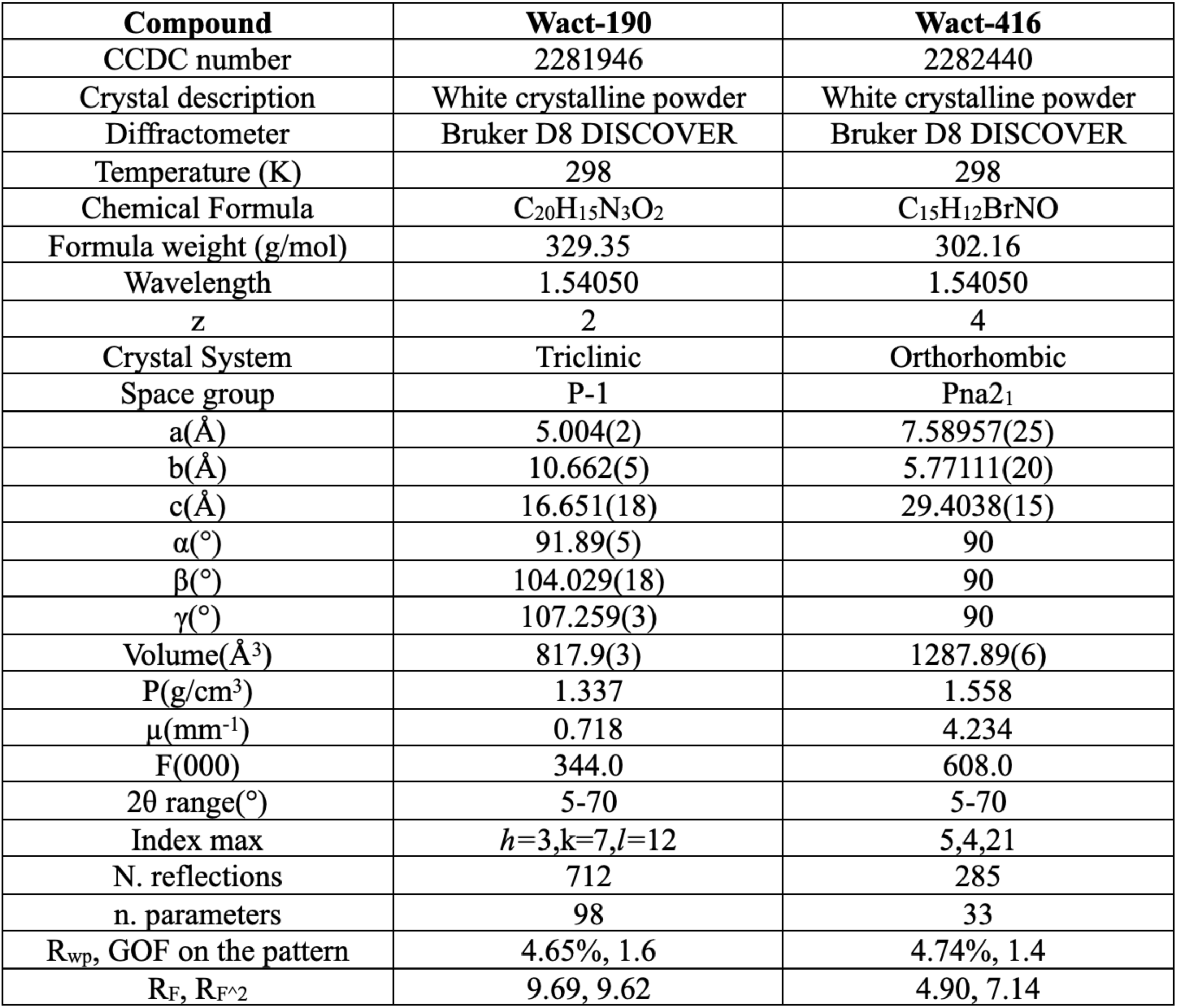
Crystallographic data collection and refinement statistics data for the crystal of wact-190 and wact-416.

**Supplemental Table 3.**
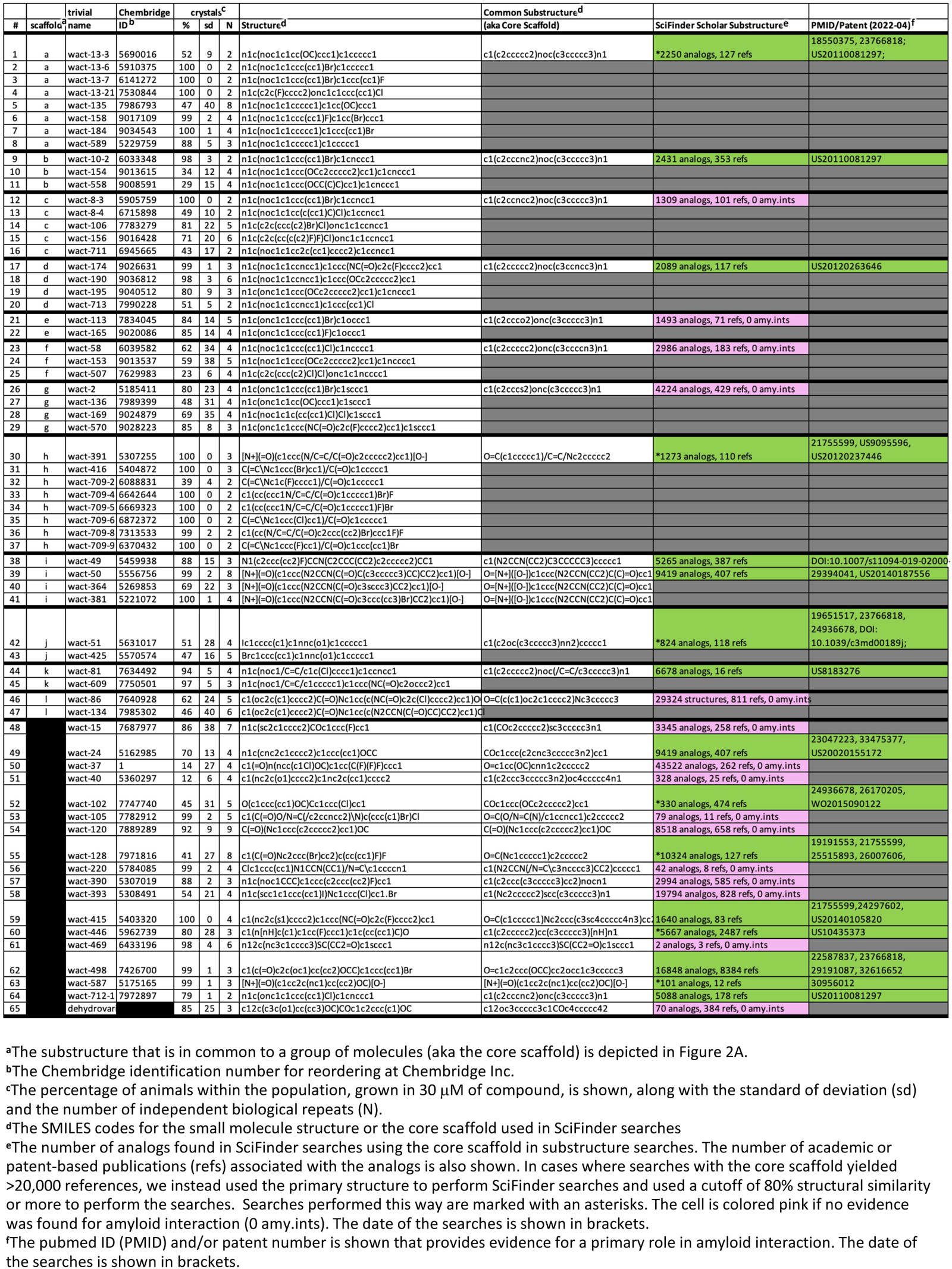
A Literature-Based Search for Evidence of Amyloid Interaction Among 65 Crystalizing Molecules.

**Supplemental Table 4.**
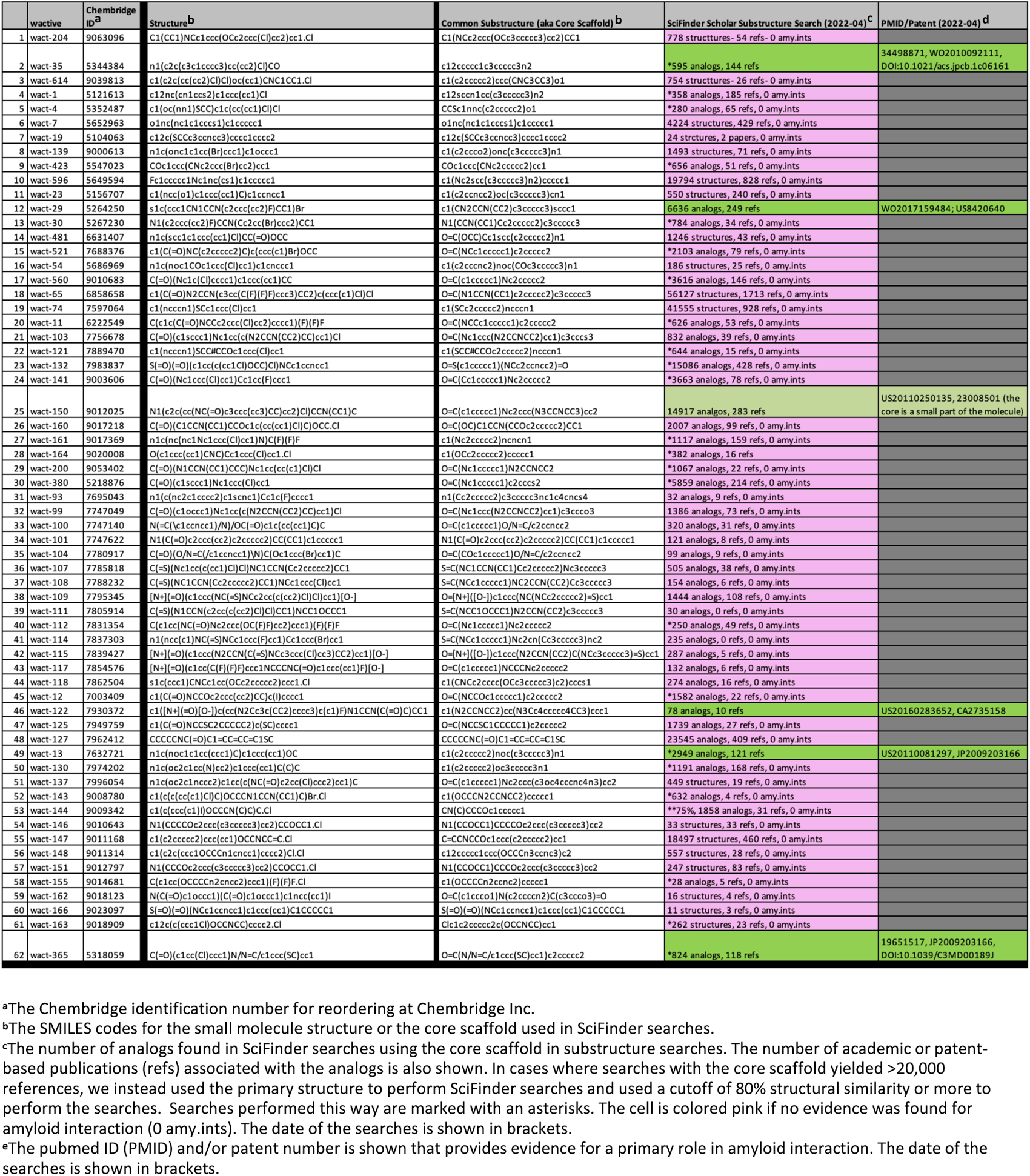
A Literature-Based Search for Evidence of Amyloid Interaction Among 62 Molecules that Fail to Form Crystals.

**Supplemental Table 5.**
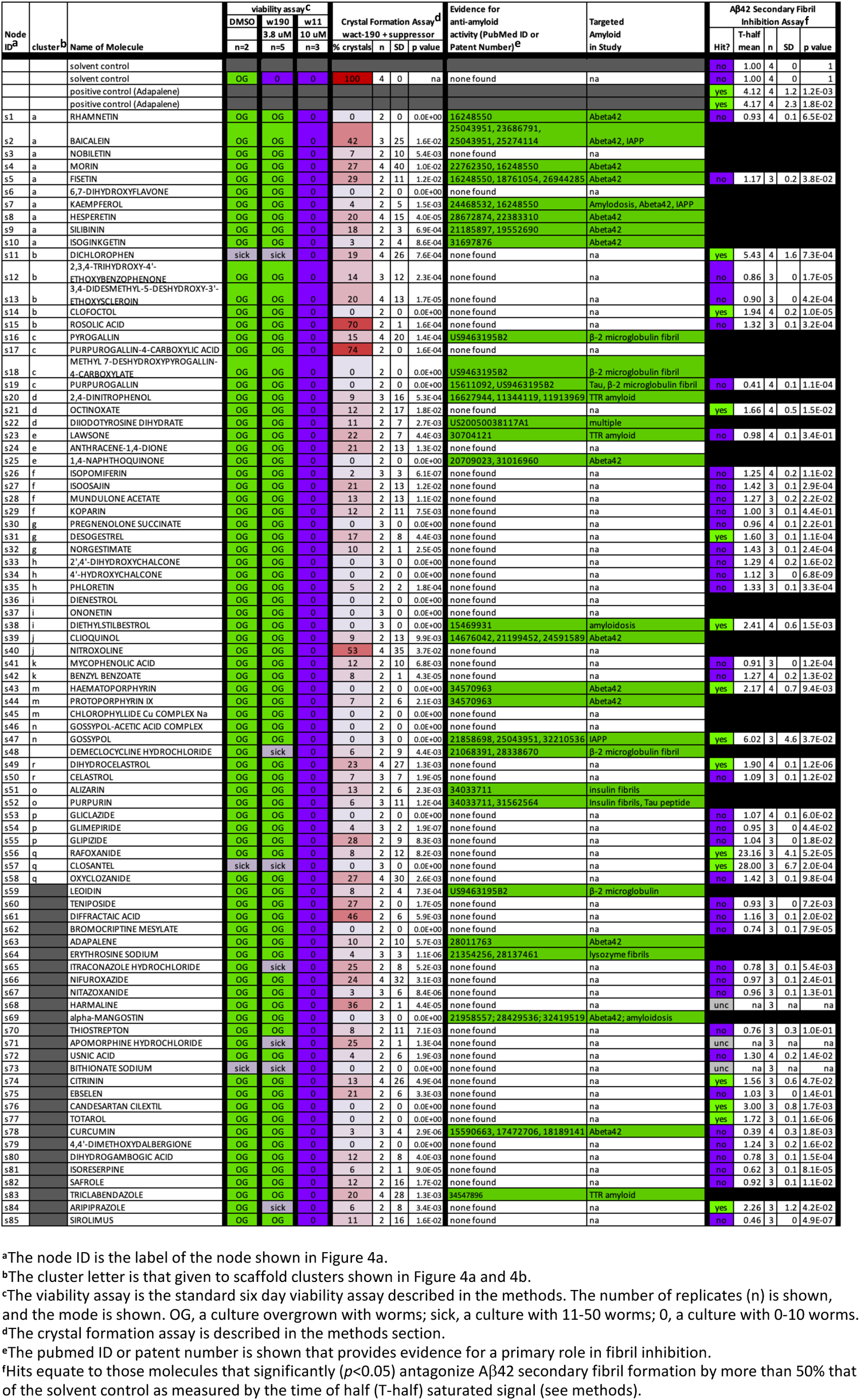
85 Spectrum Suppressors of Small Molecule Crystallization in the Pharynx Cuticle.

**Supplemental Table 6.**
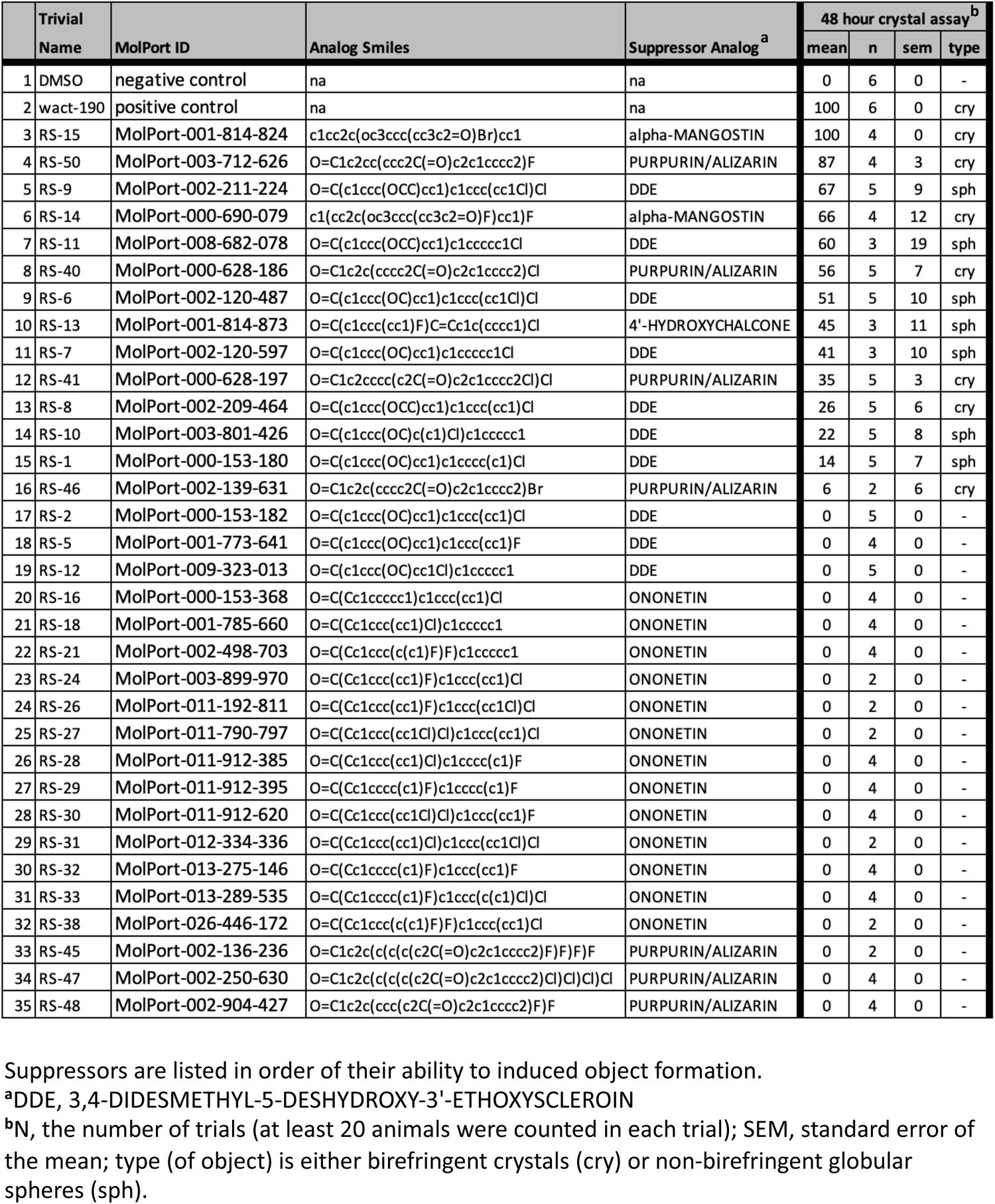
Reduced Analogs of Suppressors.

**Supplemental Table 7.**
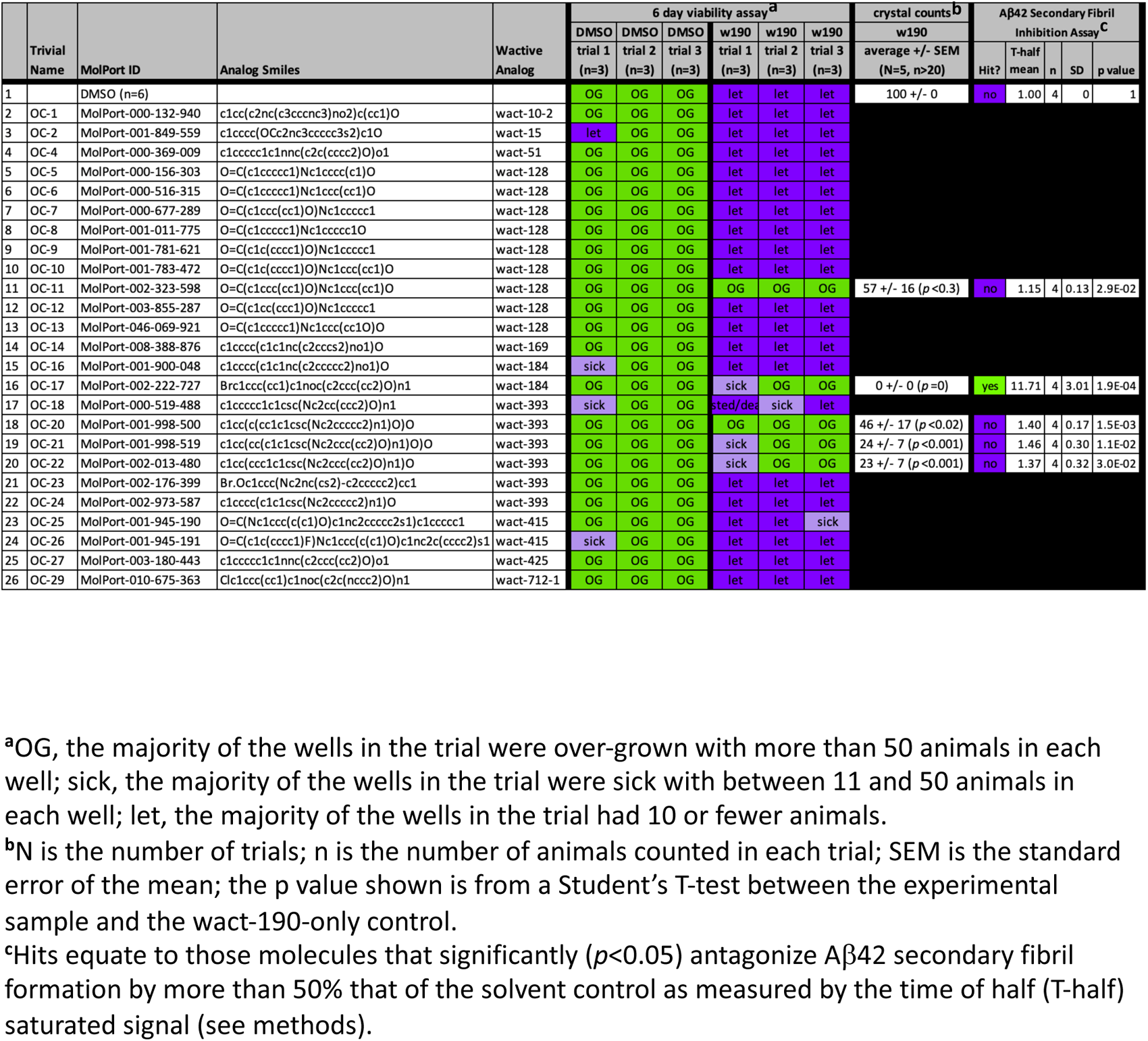
Hydroxylated Analogs of Crystal-Forming Molecules.

**Supplemental Figure 1.**
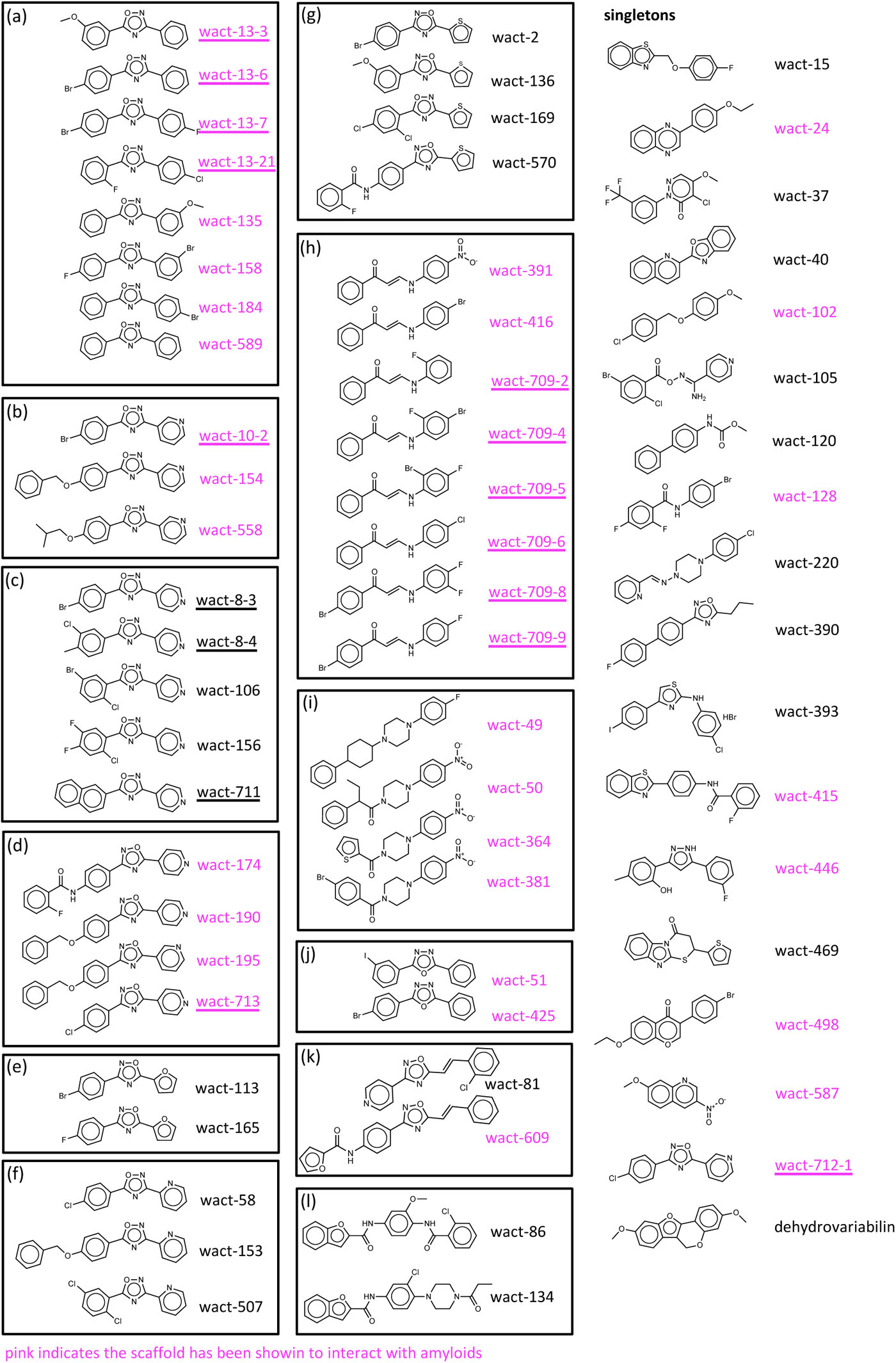
Crystallizing Small Molecules Grouped According to Structural Similarity. Small molecules belonging to structurally distinct families are boxed together. The 18 singletons are unboxed. The cluster that each boxed set of molecules belong to corresponds to the notation in Figure 3 and is indicated in the upper left-hand corner. Wactive names highlighted in pink have core scaffolds that are known to interact with amyloids (see methods and Supplemental Data File 1 for details). Molecule names that are underlined were discovered from a screen of the RLL1200 small molecule library.

**Supplemental Figure 2.**
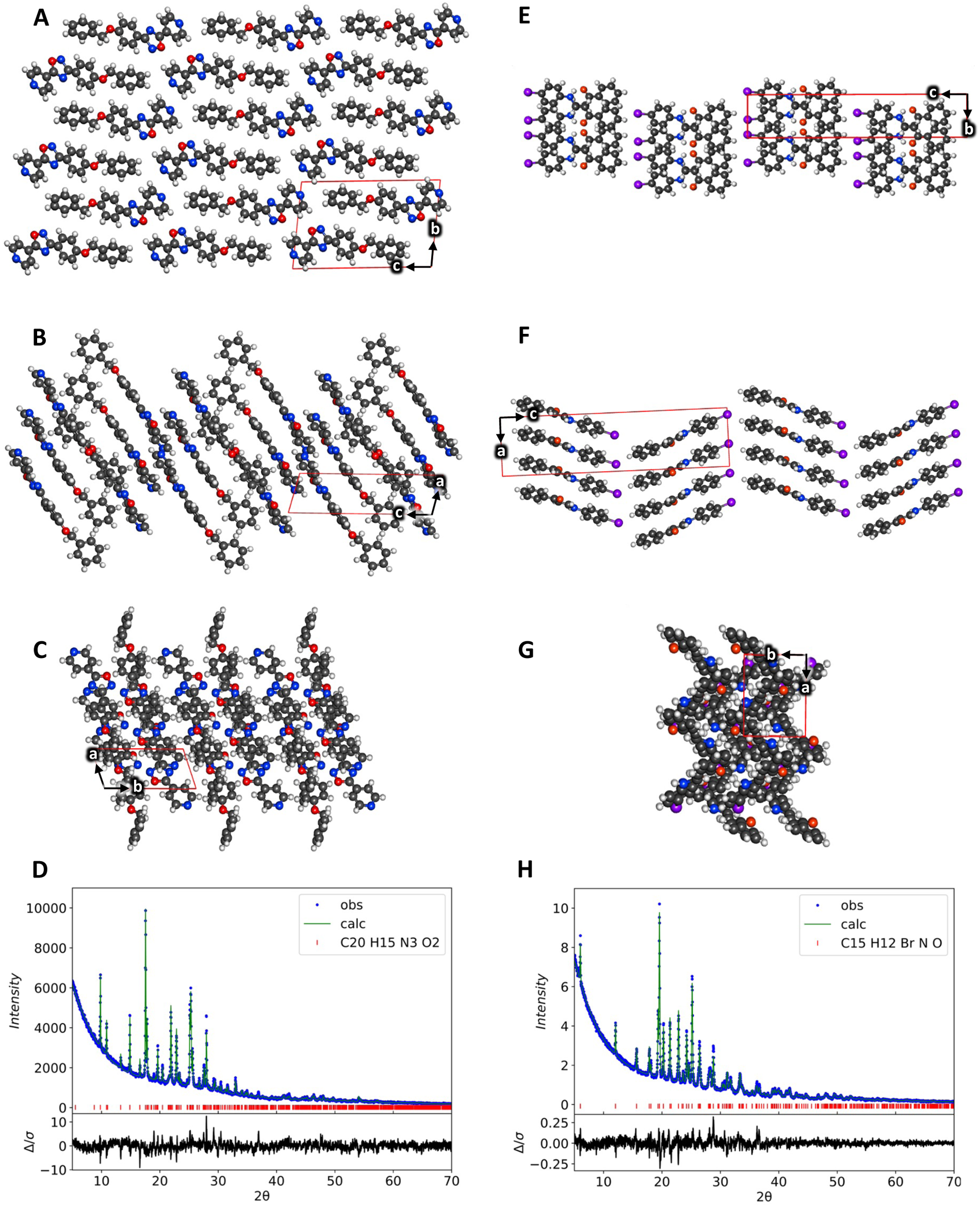
Further Details on the Biophysical Characterization of wact-190 and wact-416 *in vitro* Crystals. **A-C.** Different aspects of the 3D packing arrangement of wact-190 crystals from the a, b, and c aspects, respectively. **D.** The fitting pattern of wact-190 was obtained from the PXRD instrument. **E-G.** Different aspects of the 3D packing arrangement of wact-416 crystals from the a, b, and c aspects, respectively. **D.** The fitting pattern of wact-416 was obtained from the PXRD instrument. Panels (D) and (H) show the experimental powder diffraction profile (°), the calculate curve by using the reported structure (green line), the peak positions (|), and the normalized difference curve (I_o_-I_c_) (black curve).

**Supplemental Figure 3.**
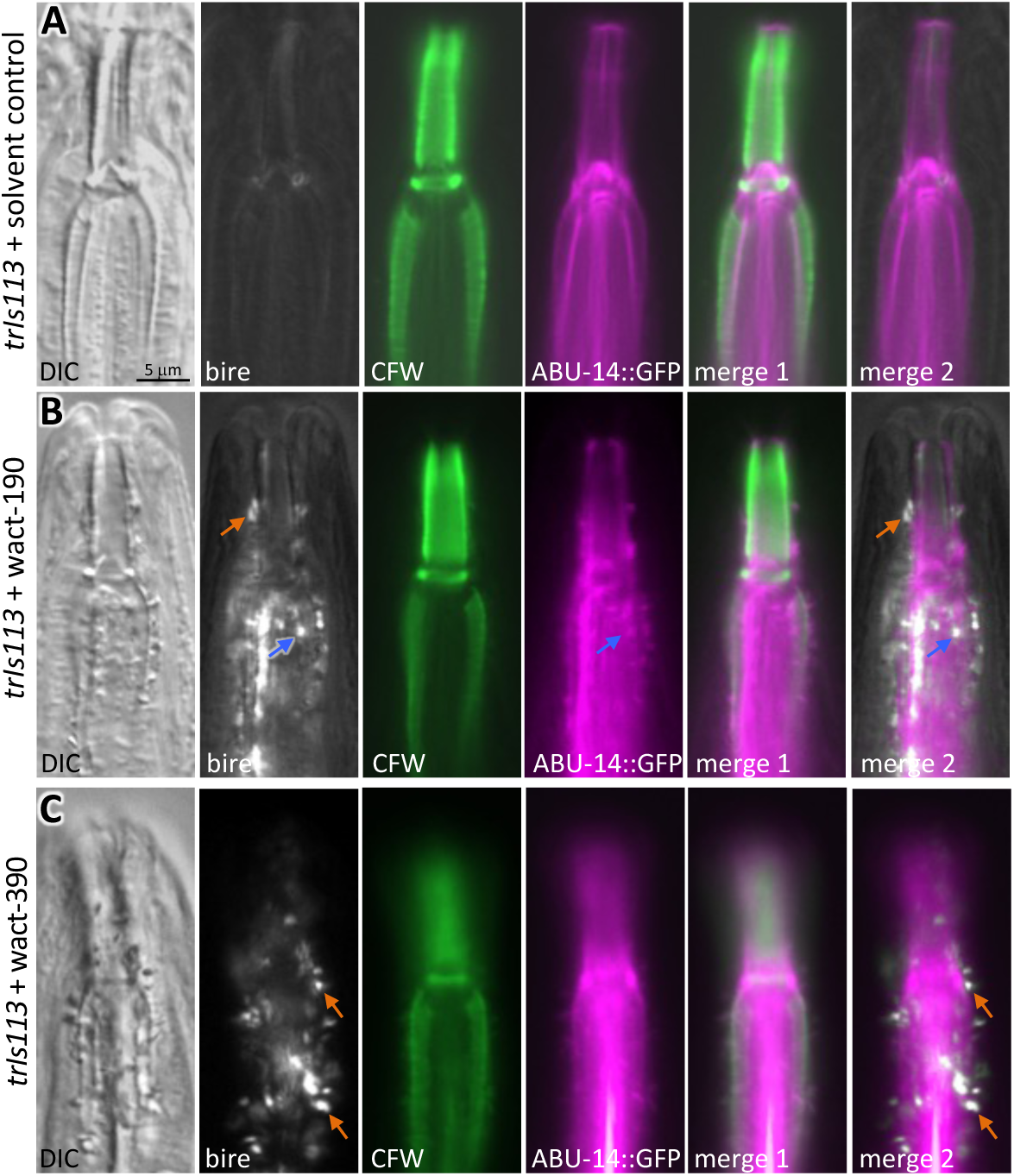
Crystal-Induced Disruption of the ABU-14::sfGFP Pharyngeal Cuticle Marker. **A-C**. Images of animals expressing ABU-14::sfGFP (the extrachromosomal array, which we integrated as *trIs113*, was a kind gift from David Raizen) and stained with the indicated dyes. The imaging channel is indicated (lower left) as is the 3-hour wactive treatment (left-hand side of micrograph series). DIC, differential interference contrast; bire, birefringence; CFW, calcofluor white. Merge 1 is the merge of ABU-14::GFP and the CFW signal. Merge 2 is the merge of the signals from ABU-14::GFP and the birefringence signal. Blue arrows indicate overlapping signals of the amyloid-binding dye and birefringence; orange arrows show a lack of amyloid-binding dye signal associated with birefringence signal. 60 μM of wact-190 and wact-390 was used in all experiments relevant to panels A-C.

**Supplemental Figure 4.**
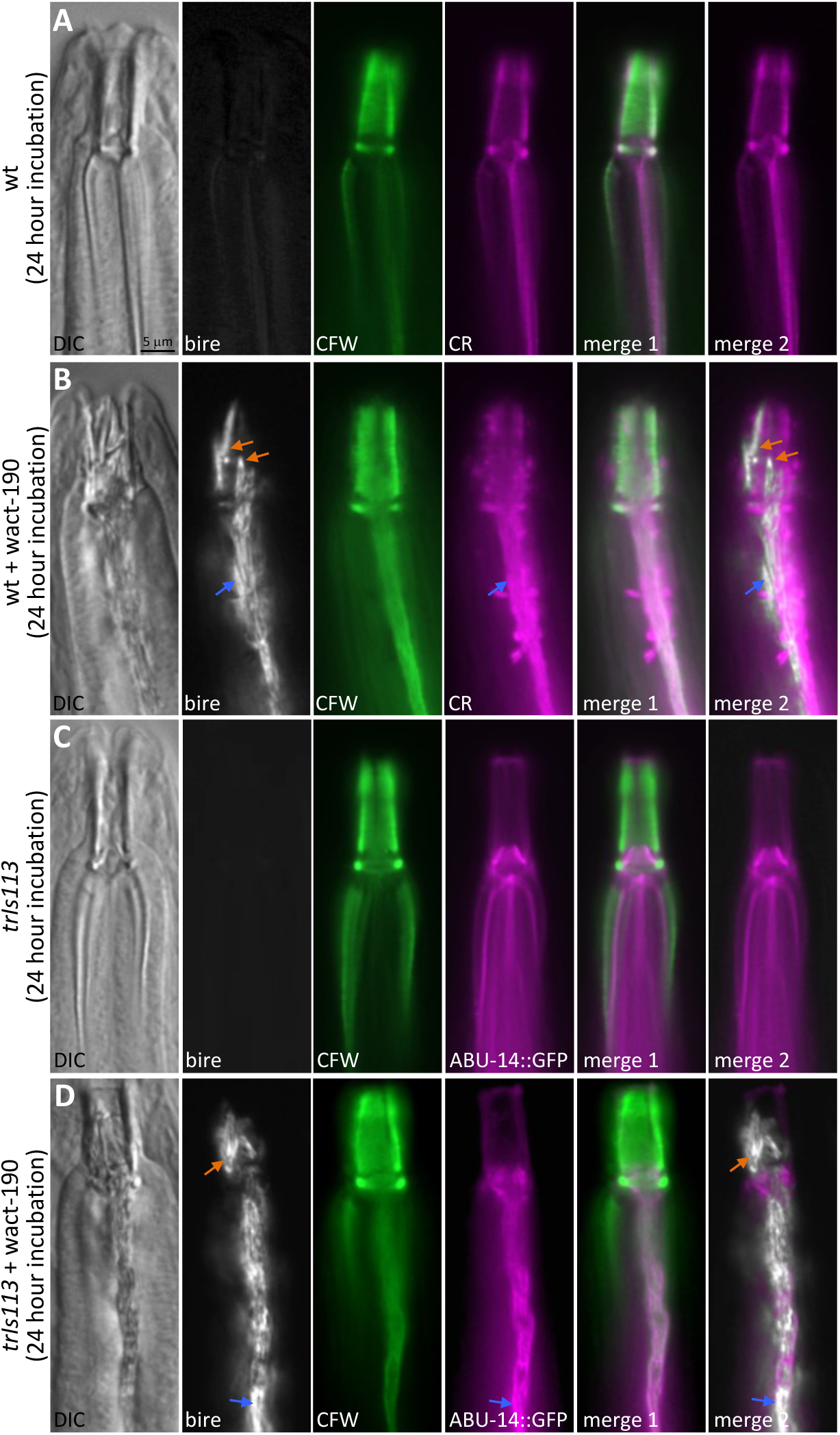
Larger wact-190 Crystals Often Fail to Overlap with Cuticle Markers. Legend is the same as that described for Figure 4 and Supplemental Figure 3.

**Supplemental Figure 5.**
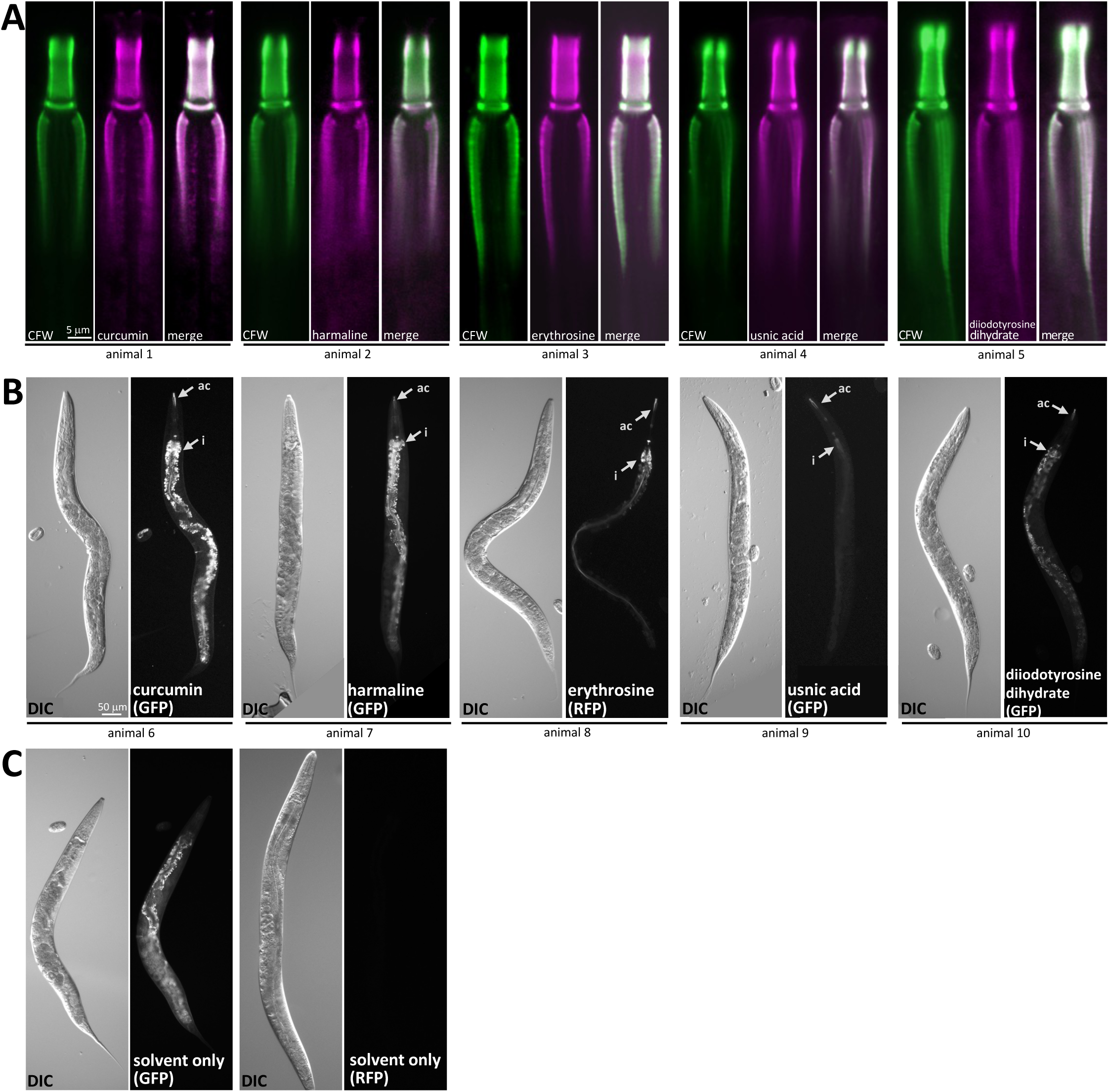
Fluorescent Suppressors Specifically Localize to the Pharyngeal Cuticle. **A.** Calcofluor white (CFW) is used as counter-stain control for each of the molecules (60 μM) examined. **B-C.** B. Whole animal pictures illustrating the specificity of the localization of the indicated suppressor molecules. The intestine exhibits autofluorescence from gut granules, as illustrated in the no-dye control pictures (C). ac, anterior corpus; i, intestine.

**Supplemental Figure 6.**
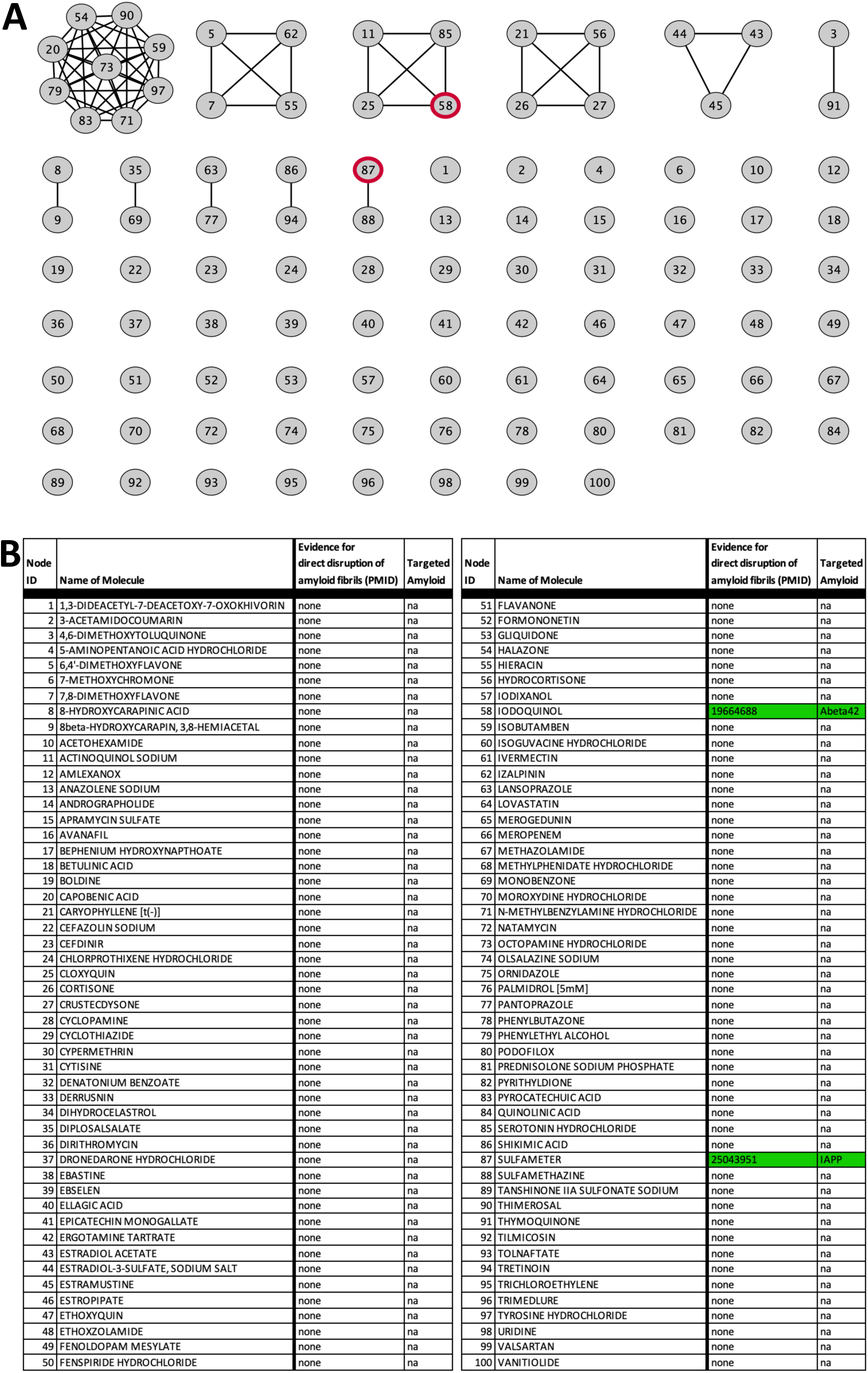
The Construction and Analysis of a Random Network from the Spectrum Library. **A.** A structural similarity network of the 100 randomly chosen Spectrum library molecules. The nodes (circles) represent a drug or natural product whose number corresponds to the identity indicated in panel B. The links (straight lines) represent a Tanimoto score of structural similarity of 0.8 or more (see methods). A bold red outline indicates that the molecule is a known disruptors of amyloid fibril formation (see panel B for details). **B.** The 100 molecules randomly chosen from the Spectrum Library that were used the build the network depicted in panel A. The node ID is the label of the node shown in panel A. The PubMed ID is shown for those molecules for which evidence could be found of a primary role in fibril inhibition.

**Supplemental Figure 7.**
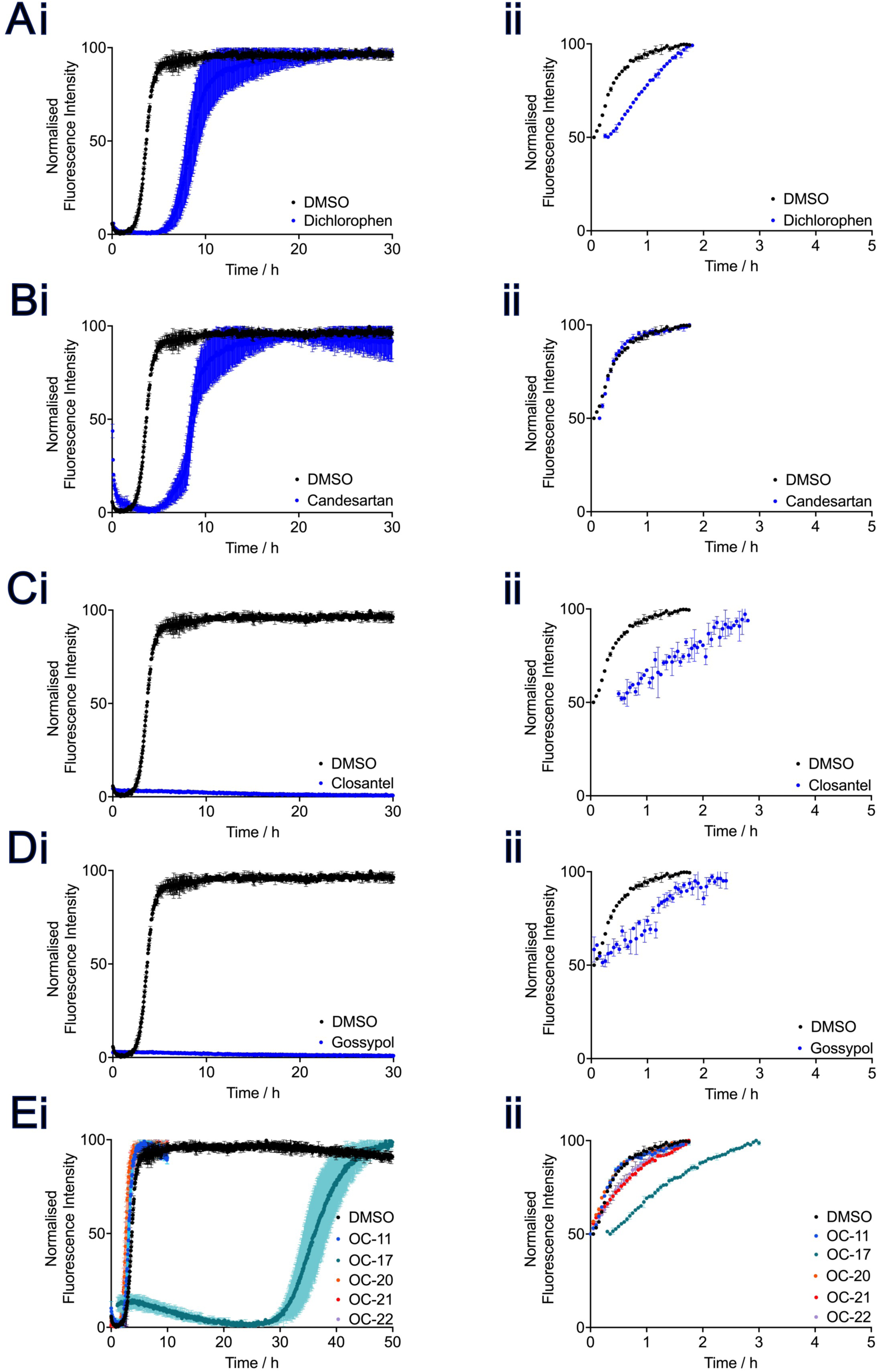
A-E. Behaviour of 2 mM Aβ42 Incubated with Small Molecules under Different Aggregation Conditions. **A.** Dichlorophen **B.** Candesartan Cilexetil **C.** Closantel **D.** Gossypol **E.** OC-11, OC-17, OC-20, OC-21, OC-22 in **i.** unseeded conditions or with **ii.** 1 mM monomer equivalent pre-formed seed (heavy seeded). Heavy seeded conditions isolate elongation as the dominant mechanism of aggregation, as the availability of fibril ends is so high that further amyloid formation only occurs via this route. By comparison, light seeded conditions (main text) are dominated by secondary nucleation and elongation, while under unseeded conditions primary nucleation, secondary nucleation and elongation are operative.

## Notes

### Competing Interest Statement

The authors have declared no competing interest.

### Summary of Updates

We revised the original version of this manuscript significantly. The original manuscript was split into two manuscripts, each of which was expanded significantly. The revised version posted here focuses on the main theme of the original version- identifying small molecules that disrupt amyloid formation.

